# Impaired OTUD7A-dependent Ankyrin regulation mediates neuronal dysfunction in mouse and human models of the 15q13.3 microdeletion syndrome

**DOI:** 10.1101/2022.01.11.475723

**Authors:** Brianna K. Unda, Leon Chalil, Sehyoun Yoon, Savannah Kilpatrick, Sansi Xing, Nadeem Murtaza, Anran Cheng, Alexandria Afonso, Elizabeth McCready, Gabriel Ronen, Jennifer Howe, Aurélie Caye-Eude, Alain Verloes, Brad W. Doble, Laurence Faivre, Antonio Vitobello, Stephen W Scherer, Yu Lu, Peter Penzes, Karun K. Singh

**Author notes:** Corresponding author: Karun K. Singh.

## Abstract

Copy number variations (CNV) are associated with psychiatric and neurodevelopmental disorders (NDDs), and most, including the recurrent 15q13.3 microdeletion disorder, have unknown disease mechanisms. We used a heterozygous 15q13.3 microdeletion mouse model and patient iPSC-derived neurons to reveal developmental defects in neuronal maturation and network activity. To identify the underlying molecular dysfunction, we developed a neuron-specific proximity-labeling proteomics (BioID2) pipeline, combined with patient mutations, to target the 15q13.3 CNV genetic driver *OTUD7A*. *OTUD7A* is an emerging independent NDD risk gene with no known function in the brain, but has putative deubiquitinase (DUB) function. The OTUD7A protein-protein interaction (PPI) network revealed interactions with synaptic, axonal, and cytoskeletal proteins and was enriched for known ASD and epilepsy risk genes. The interactions between OTUD7A and the NDD risk genes Ankyrin-G (*Ank3*) and Ankyrin-B (*Ank2*) were disrupted by an epilepsy-associated *OTUD7A* L233F variant. Further investigation of Ankyrin-G in mouse and human 15q13.3 microdeletion and OTUD7A*^L233F/L233F^* models revealed protein instability, increased polyubiquitination, and decreased levels in the axon initial segment (AIS), while structured illumination microscopy identified reduced Ankyrin-G nanodomains in dendritic spines. Functional analysis of human 15q13.3 microdeletion and OTUD7A*^L233F/L233F^* models revealed shared and distinct impairments to axonal growth and intrinsic excitability. Importantly, restoring OTUD7A or Ankyrin-G expression in 15q13.3 microdeletion neurons led to a reversal of abnormalities. These data reveal a critical OTUD7A-Ankyrin pathway in neuronal development, which is impaired in the 15q13.3 microdeletion syndrome, leading to neuronal dysfunction. Further, our study highlights the utility of targeting CNV genes using cell-type specific proteomics to identify shared and unexplored disease mechanisms across NDDs.

## INTRODUCTION

Genomic copy number variations (CNVs) are structural variations that involve deletions or duplications of segments of DNA. They are frequently associated with disease, and represent a major class of genetic risk factors for neurodevelopmental disorders (NDDs), with large recurrent deletions causing the most severe outcomes ^1, 2^. Due to a lack of understanding of disease mechanisms, there are no specific treatments for NDDs caused by CNVs. Evidence indicates that the loss/gain of certain genes within a CNV, known as ‘genetic drivers’, are the major cause of neurological deficits^3–9^; however, genetic mechanisms outside of the CNV region, as well as pleiotropic and polygenic contributions, also play a role^10–12^. Given that multiple genes are located within a CNV, studies have focused on genetic drivers to identify relevant disease mechanisms and molecular targets for potential intervention. Examples include the 22q11.2, 16p11.2, and 7q23.11 deletion syndromes where gene(s) within the CNV are responsible for specific developmental or neuronal deficits, and in some cases, the identification of genetic drivers has allowed the development of rescue strategies^3–9^.

The 15q13.3 1.53 Mb microdeletion syndrome (MIM: 612001) locus (chr15:30,910,306–32,445,407 [hg19]) that resides within breakpoints BP4-BP5 on human chromosome 15 is a recurrent CNV that displays incomplete penetrance and expressivity. It is strongly associated with a heterogeneous set of phenotypes including intellectual disability (ID), autism spectrum disorder (ASD), epilepsy (MIM: 607208), and schizophrenia^13–21^. Individuals are typically heterozygous for the 15q13.3 microdeletion, which encompasses seven protein-coding genes, one microRNA, and two putative pseudogenes (*ARHGAP11BI* [MIM: 616310], *LOC100288637*, *FAN1* [MIM: 613534], *MTMR10*, *TRPM1* [MIM: 603576], *LOC283710*, microRNA-211, *KLF13* [MIM: 605328], *OTUD7A* [MIM: 612024], and *CHRNA7* [MIM: 118511])^22^. Mouse models of the 15q13.3 deletion display cortical dysfunction, behavioral abnormalities, and epilepsy, which are consistent with a developmental etiology^23–27^. Cortical excitatory and inhibitory neurons have both been implicated ^26, 28–30^, but how dysfunction of either of these cell types occurs at the molecular level is not understood. Human studies on 15q13.3 microdeletion iPSCs have identified transcriptional alterations in developing cortical neural cells, but the functional consequences remain unknow^31^. Previous work has identified several candidate genes in the locus contributing to the clinical phenotype including Cholinergic Receptor Nicotinic Alpha 7 Subunit (*CHRNA7*), Fanconi-associated nuclease 1 (*FAN1*) and Ovarian tumor domain containing protein 7A (*OTUD7A*)^18, 21, 30, 32–36^. However, it remains unknown how loss of candidate driver genes in the 15q13.3 deletion leads to specific deficits and molecular abnormalities in the developing brain.

Recent clinical findings suggest that among the potential contributing genes in the 15q13.3 microdeletion, *OTUD7A* (a putative deubiquitinase) may play a prominent role. Case reports revealed that homozygous loss of function mutations in *OTUD7A* are associated with severe epilepsy, dystonia and intellectual disability^37, 38^. Additionally, emerging genetic sequencing studies have provided new evidence that supports OTUD7A as an independent NDD risk gene^39–41^. This is consistent with *OTUD7A* having the highest intolerance score (pLI) for loss-of-function alleles within the 15q13.3 locus^30^, indicating that mutations could provide clues to the function of OTUD7A. Additionally, studies using heterozygous 15q13.3 microdeletion and *Otud7a* knockout (KO) mouse models have demonstrated a strong role for *OTUD7A* in the regulation of cortical excitatory neuron development^30, 36^. At the molecular level, OTUD7A has been linked to TRAF6 as an interacting partner in non-neuronal cells^42^, but the proteins and pathways that OTUD7A interacts with or regulates in the brain remain unknown.

To understand the cellular mechanisms involved in the 15q13.3 microdeletion, we utilized the 15q13.3 microdeletion mouse model [*Df(h15q13)/+*] and human induced pluripotent stem cell (iPSC)-derived induced glutamatergic neurons (iNeurons) from three unrelated 15q13.3 microdeletion probands and their familial controls. We also analyzed iNeurons from a proband with epileptic encephalopathy and a homozygous missense variant in *OTUD7A* [NM_130901.2:c.697C>T, p.(Leu233Phe)] to gain further insight into the importance of *OTUD7A*^37^. In mouse *Df(h15q13)/+* and human 15q13.3 microdeletion/ OTUD7A^L233F/L233F^ models, we identified common impairments in dendrite and dendritic spine morphology, as well as population level spontaneous firing using long-term multielectrode arrays (MEAs) (Fig. 1a). Mapping of CNV driver gene signaling networks is critical to understanding disease mechanisms but poses a challenge because many pathways can be involved. We turned to proteomic tools to navigate such complex questions^8, 43–51^. Traditional proteomic screens can generate an abundance of data, making it difficult to narrow down the critical signaling mechanisms. To address this, we developed a lentiviral neuron-specific proximity-labeling proteomics strategy (BioID2) to identify the protein-protein interaction (PPI) network of OTUD7A and compared this to independent *OTUD7A* mutations linked to ASD (*OTUD7A*-N492_K494del) and epilepsy (*OTUD7A* L233F). We discovered that OTUD7A is associated with synaptic, axonal, cytoskeletal, and NDD risk gene networks, which were differentially disrupted by the mutations. We confirmed the interaction of OTUD7A with the NDD risk genes Ankyrin-G (*Ank3*) and Ankyrin-B (*Ank2*), which regulate different aspects of the growth and structure of dendritic spines, axon initial segment (AIS) and axons^52–58^. Further analysis of Ankyrin-G revealed reduced levels in dendritic spines and the AIS in 15q13.3 microdeletion and OTUD7A^L233F/L233F^ iNeurons, as well as reduced protein stability and elevated ubiquitination. We identified novel deficits in axon growth and intrinsic excitability in 15q13.3 microdeletion iNeurons, further supporting the importance of the AIS/axonal OTUD7A PPI network. Importantly, the neuronal impairments in *Df(h15q13)/+* mouse cortical neurons and patient iNeurons were rescued upon restoration of OTUD7A or Ankyrin-G expression (Fig. 1b). Our collective data suggest that the regulation of Ankyrin protein homeostasis by OTUD7A is a critical mechanism for neurodevelopment, which is impaired in the 15q13.3 microdeletion. This also provides the basis for potential therapeutic strategies to reverse neuronal deficits.

**Figure 1.**
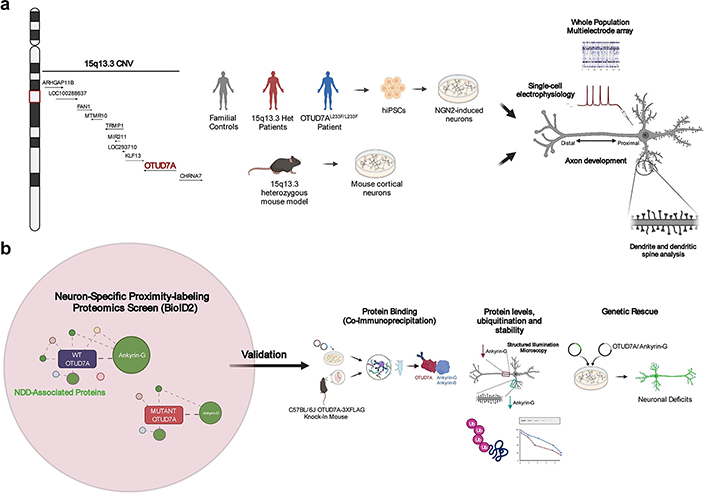
Overview of study workflow. **(a)** 15q13.3 microdeletion locus and driver gene, *OTUD7A* (red font). Mouse and human15q13.3 microdeletion and *OTUD7A* patient mutation models show abnormal neuronal morphology and electrical activity. **(b)** A BioID2 proximity-labeling proteomics screen of OTUD7A identified enrichment of proteins localized to the postsynaptic density, cytoskeleton and axon, which are disrupted by *OTUD7A* mutations. Additionally, the OTUD7A PPI network was enriched for known NDD-associated proteins. Top BioID2 interactors (including Ankyrin-G) were validated via co-immunoprecipitation, protein levels, ubiquitination status and protein stability in mouse and human models, and genetic rescue of morphological abnormalities in the 15q13.3 microdeletion background.

## RESULTS

### *Df(h15q13)/+* cortical neurons show developmental impairment of population-level spontaneous firing and bursting patterns

We previously reported that *Df(h15q13)/+* mouse cortical neurons display a decrease in dendritic spine density and a shift to a reduced proportion of mushroom spines and a higher proportion of stubby type spines, suggesting that *Df(h15q13)/+* neurons may be immature^30^.To investigate if there is a functional consequence, we used *in vitro* multi-electrode arrays (MEAs) to record live spontaneous neuronal activity over time from WT and *Df(h15q13)/+* cortical neurons. We measured spikes (action potentials) from cultured cortical neurons and analyzed firing rate, bursting activity, and network bursting. At days-in-vitro (DIV) 7*, Df(h15q13)/+* neurons displayed a significant decrease in active electrodes (**Supplementary Figure 1a**), which disappeared from DIV 14 onward, suggesting an early defect in activity. *Df(h15q13)/+* neurons showed a significant decrease in weighted mean firing rate (wMFR), which considers the number of active electrodes, from DIV 14 onwards (Fig. 2a,b). *Df(h15q13)/+* cortical neurons also showed a significant decrease in the frequency of bursts (**Supplementary Figure 1b**) and network bursts (Fig. 2a,c) from DIV 21 onward. These results suggest that *Df(h15q13)/+* neurons develop reduced levels of spontaneous activity, as well as reduced levels of coordinated activity between neurons within a well, which persist up to approximately 4 weeks in culture.

**Figure 2.**
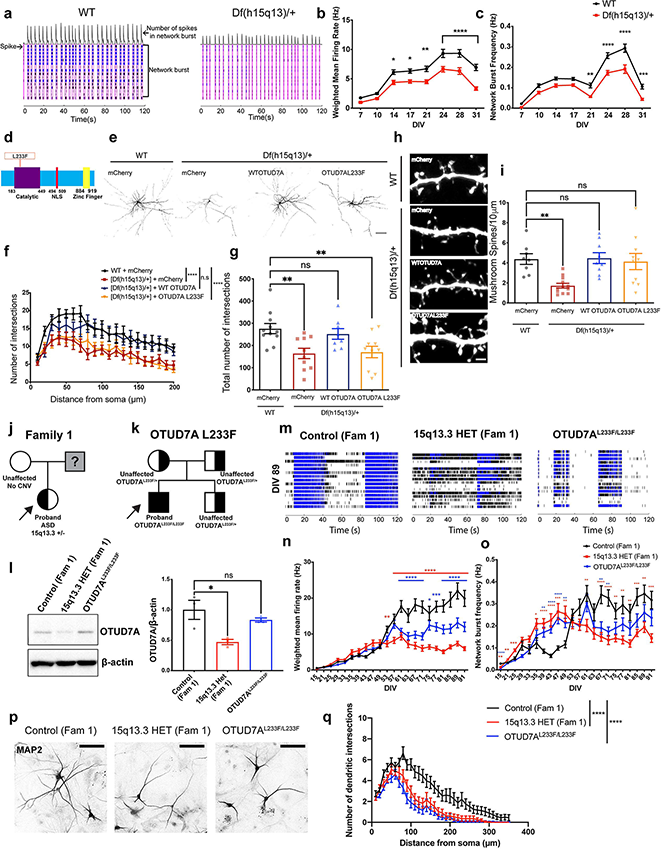
*Df(h15q13)/+* cortical neurons and human patient iNeurons show impairments in synaptic morphology and persistent functional deficits. **(a)** Raster plots of MEA recordings of neural network activity from DIV 24 WT and *Df(h15q13)/+* mouse cortical neurons. n=77 wells WT, 65 wells *Df(h15q13)/+* from 3 mouse cortical cultures on 3 MEA plates. **(b)** Weighted mean firing rate. Repeated Measures 2-Way ANOVA with Sidak’s post-hoc test, *p<0.05, **p<0.01, ****p<0.0001; Interaction: F(7,980)=3.950, p=0.0003; DIV: F(7,980)=87.98, p<0.0001; Genotype: F(1,140)=30.64, p<0.0001; Subject: F(140,980)=4.436, p<0.0001. **(c)** Network burst frequency. Repeated Measures 2-Way ANOVA with Sidak’s post-hoc test, **p<0.01, ***p<0.001, ****p<0.0001; Interaction: F(7,980)=4.341, p<0.0001; DIV: F(7,980)=115.2, p<0.0001; Genotype: F(1,140)=40.27; Subject: F(140,980)=2.78, p<0.0001. **(d)** Schematic of the human OTUD7A protein showing the location of the L233F variant. **(e)** Representative confocal images from co-transfected DIV 14 WT and *Df(h15q13)/+* mouse cortical neurons; 20X objective, scale bar 100 μm. **(f-g)** Sholl analysis. n= 11 neurons WT + mCherry, 10 neurons [Df(h15q13)/+] + mCherry, 8 neurons [Df(h15q13)/+] + WT OTUD7A-mCherry, 10 neurons [Df(h15q13)/+] + OTUD7A L233F. Samples were taken from 3 mouse cultures. **(f)** 2-Way ANOVA with Dunnett’s post-hoc test; ****p<0.0001; Interaction: F(57,700)=0.505, p=0.9991; DIV: F(19,700)=10.54, p<0.0001; Distance: F(3,700)=57.12, 0<0.0001 **(g)** Total number of dendritic intersections. One-Way ANOVA with Dunnett’s post-hoc test, **p<0.01, F(3,35)=5.775, p=0.0026. **(h)** Representative confocal images of dendritic segments from co-transfected DIV 14 WT and *Df(h15q13)/+* mouse cortical neurons; 63X objective, scale bar 2 μm. **(i)** Mushroom spine density. n= 8 neurons WT + mCherry, 12 neurons [Df(h15q13)/+] + mCherry, 9 neurons [Df(h15q13)/+] + WT OTUD7A-mCherry, 10 neurons [Df(h15q13)/+] + OTUD7A L233F. Samples were taken from 3 mouse cultures. *p<0.05, **p<0.01; One-Way ANOVA with Dunnett’s post-hoc test; F(3,35)= 6.423, p=0.0014. **(j)** Pedigree of Family 1 and **(k)** the OTUD7A L233F Family. **(l)** Representative western blot (left) and quantification (right) of OTUD7A levels in iNeurons; n= 3 separate Ngn2/Rtta transductions per line; One-Way ANOVA with Dunnett’s post-hoc test, *p<0.05, F(2.6)=7.921, p=0.0207. **(m)** Raster plots of MEA recordings of neural network activity from DIV 89 Family 1 and OTUD7A^L233F/L233F^ human iNeurons. Control (Fam 1) n=29 wells, 15q13.3 HET (Fam 1) n=29 wells, OTUD7A^L233F/L233F^ n=30 wells from two separate NGN2/Rtta transductions. **(n)** Weighted mean firing rate. Repeated measures Two-Way ANOVA with Dunnett’s post-hoc test; Interaction: F(42,1785)=17.59, p<0.0001; DIV: F(2.925, 248.6)=108.2, p<0.0001; Genotype: F(2.85)=15.96, p<0.0001; Subject: F(85,1785)=16.12, p<0.0001. **(o)** Network burst frequency. Repeated measures Two-Way ANOVA with Dunnett’s post-hoc test; *p<0.05, **p<0.01, ***p<0.001, ****p<0.0001. **(p)** Representative images and **(q)** sholl analysis of MAP2-positive Family 1 and OTUD7A^L233F/L233F^ proband human iNeurons; 20X objective, scale bar 100 μm. Control n= 33 neurons, 15q13.3 HET n= 31 neurons, OTUD7A^L233F/L233F^ n=38; ***p<0.001, 2-Way ANOVA with Dunnett’s post-hoc test; Interaction: F(68,3465)=1.943, p<0.0001; Distance from soma: F(34,3465)=53.30, p<0.0001; Genotype: F(2, 3465) = 200.2, p<0.0001.

### An epileptic encephalopathy-associated OTUD7A L233F variant disrupts dendrite development in cortical neurons

The first case of a homozygous missense variant in *OTUD7A* chr15:g.31819467G>A [NM_130901.2:c.697C>T, p.(Leu233Phe)], hereafter referred to as OTUD7A L233F, was reported in a two-year-old male with severe global developmental delay, language impairment and epileptic encephalopathy^37^. This variant was inherited from both parents, each heterozygous and displaying mild learning difficulties^37^ (**Supplementary Figure 2a;** Fig. 2k).The variant results in a substitution of leucine 233 to phenylalanine within the catalytic domain of OTUD7A, suggesting it may impact protein function in neurons (Fig. 2d). We examined the impact of the OTUD7A L233F variant by testing whether it could rescue the decrease in dendrite complexity and mushroom spine density in *Df(h15q13)/+* neurons. We created mCherry-tagged WT OTUD7A and OTUD7A L233F expression constructs (**Fig. S1 c**), which we co-expressed with Venus (for visualization) in WT and *Df(h15q13)/+* neurons. Fluorescence intensities of WT OTUD7A-mCherry and OTUD7A L233F-mCherry in neurons were the same (**Supplementary Figure 1d-f**), suggesting that the variant does not change OTUD7A protein levels. Sholl analysis revealed that WT OTUD7A rescued dendrite complexity in *Df(h15q13)/+* neurons, whereas OTUD7A L233F did not, indicating that the OTUD7A L233F variant affects the ability of OTUD7A to regulate dendrite complexity (Fig 2e-g). Next, we examined dendritic spine development in *Df(h15q13)/+* neurons and found that spine density was decreased, which was rescued by both WT OTUD7A and OTUD7A L233F (**Supplementary Figure 1g**). Further morphological analysis showed a decrease in mushroom spine density, which was also rescued by both WT OTUD7A and OTUD7A L233F (Fig. 2h,i), and no change in stubby spine density (**Supplementary Figure 1h**). *Df(h15q13)/+* neurons also displayed a decrease in the proportion of spines classified as mushroom type spines, which was again rescued by both WT OTUD7A and OTUD7A L233F (**Supplementary Figure 1i**), These results suggest that the predominant impact of the OTUD7A L233F variant is on dendritic arborization.

### 15q13.3 microdeletion and OTUD7A L233F patient iNeurons display impairments in spontaneous firing and dendritic arborization

Given the synaptic activity and morphological alterations in mouse *Df(h15q13)/+* cortical neurons, we next examined whether these deficits would be recapitulated in 15q13.3 microdeletion iPSC-derived iNeurons. We generated iPSCs from peripheral mononuclear blood cells obtained from three unrelated individuals with a 15q13.3 deletion and available familial controls, as well as the individual with a homozygous missense variant (L233F) in *OTUD7A* (**Supplementary Figure 2a,b**). iPSCs were generated using Sendai virus, and all lines had a normal karyotype, were mycoplasma negative and expressed pluripotency markers (**Supplementary Figure 3**). The first family (Family 1) consists of a female ASD proband with a 15q13.3 deletion (BP4-5) and the proband’s neurotypical mother, who does not have a 15q13.3 deletion (Fig. 2j**; Supplementary Figure 2a,b)**. The second proband (Family 2) is a female with a 15q13.3 deletion (BP4-5) along with absence seizures, developmental delay (DD), intellectual disability (ID), and a learning disorder. Family 2 also displays incomplete penetrance, as the proband’s mother is neurotypical but carries the deletion (**Supplementary Figure 2a,b; Supplementary Figure 4d**). The proband’s neurotypical brother (no 15q13.3 deletion) was used as a familial control (**Supplementary Figure 2a,b; Supplementary Figure 4d**). The third proband (Family 3) has a15q13.3 deletion (BP4-5) and has ASD, attention deficit hyperactivity disorder (ADHD) and up to 150 absence seizures per day. The neurotypical father (no 15q13.3 deletion) is the familial control (**Supplementary Figure 2a,b; Supplementary Figure 4f**). The deletion in all probands and carrier encompasses *FAN1*, *MTMR10*, *TRPM1*, *MIR211*, *KLF13*, *OTUD7A* and *CHRNA7*. However, *ARHGAP11B* is deleted in the family 1 and 3 deletions, but not in the family 2 deletion. iPSCs were also generated from the independent proband with the homozygous *OTUD7A* variant (OTUD7A^L233F/L233F^) (Fig. 2k**; Supplementary Figure 2a).** Since both parents and brother carry the deletion, a familial control was not available. Therefore, experiments on this proband were performed in conjunction with the 15q13.3 microdeletion proband and familial control from Family 1.

Considering the observed defects in cortical excitatory neurons from *Df(h15q13)/+* mice in this study and our previous research^30^, we generated human glutamatergic neurons (iNeurons) using ectopic expression of Neurogenin-2 (*NGN2*) in iPSCs, which are used extensively to model neurodevelopmental disorders^65–71^, allowing us to test whether there are cell-autonomous defects caused by the 15q13.3 microdeletion. iNeurons may be more heterogeneous than initially thought depending on NGN2 expression levels^72^; therefore, NGN2 and Rtta lentiviruses were titered in iPSCs to consistently achieve >90% transduction efficiency across all transductions. We confirmed that expression of a subset of genes in the 15q13.3 interval (*OTUD7A*, *MTMR10*, *FAN1* and *CHRNA7*) are reduced by ∼50% in iNeurons derived from individuals with a 15q13.3 microdeletion from all three families at 7 days post-NGN2 induction (PNI), whereas the expression of the deletion-flanking gene *TJP1* was unchanged (**Supplementary Figure 4a-c)**. OTUD7A protein levels in PNI Day 7 iNeurons were significantly reduced by ∼50% in the Family 1 and Family 2 probands, but not in the OTUD7A^L233F/L233F^ patient or in the Family 3 proband (Fig. 2l**, Supplementary Figure 4e,g**).

We examined neural network activity of iNeurons using long-term MEAs (up to 13 weeks) co-cultured on a layer of mouse glial cells. iNeurons from all three probands, the Family 2 maternal carrier, and the OTUD7A^L233F/L233F^ proband displayed an early decrease in the number of active electrodes per well, which subsequently disappeared and was maintained at control levels for the remainder of the recording window (**Supplementary Figure 5a,e,k; Supplementary Table 1**). In Family 1, the 15q13.3 and OTUD7A^L233F/L233F^ probands had reduced wMFR from approximately week 8 (DIV 53) onwards (Fig. 2m,n**; Supplementary Table 1**). Network burst frequency and burst frequency were elevated in patient neurons compared to the control; however, this phenotype reversed at approximately week 8 and the 15q13.3 and OTUD7A^L233F/L233F^ proband iNeurons had reduced bursting and network bursting for the remainder of the experiment (Fig. 2m,o**; Supplementary Figure 5b; Supplementary Table 1**). Analysis of the proband from Family 2 also revealed a reduction in weighted mean firing rate; however, this difference occurred later, at approximately 12 weeks in culture (DIV89) (**Supplementary Figure 5f; Supplementary Table 1**). Similar to the Family 1 proband, the Family 2 15q13.3 proband also displayed an initial increase in burst frequency and network burst frequency, followed by a reversal at a later timepoint (∼12 weeks) (**Supplementary Figure 5g,h; Supplementary Table 1**). The maternal carrier showed a similar reversal in network burst frequency, but did not display a similar wMFR phenotype (**Supplementary Figure 5 f,g; Supplementary Table 1**). The Family 3 proband displayed a less pronounced phenotype than the Family 1 and Family 2 probands, compared to their respective controls. wMFR, network burst frequency and burst frequency were all acutely decreased in the proband at approximately 4 weeks in culture (**Supplementary Figure 5l,m,n; Supplementary Table 1**). Interestingly, all three probands and the OTUD7A^L233F/L233F^ proband showed a decrease in the number of spikes per burst, albeit at different days in culture (**Supplementary Figure 5c,i,o; Supplementary Table 1**). The data from these experiments suggest that there are early changes in neuronal maturation as well as later changes in spontaneous neural network activity in both 15q13.3 microdeletion and OTUD7A^L233F/L233F^ patient iNeurons.

We next examined neuronal morphology in human iNeurons by measuring dendritic arborization at PNI day 28. Sholl analysis of MAP2-positive iNeurons revealed a significant reduction in dendritic arborization in the 15q13.3 microdeletion and OTUD7A^L233F/L233F^ probands (Fig. 2p,q**; Supplementary Table 1**). This morphological phenotype, combined with the impairment in spontaneous neuronal activity, resembles the morphological and functional abnormalities observed in *Df(h15q13)/+* cortical neurons. Additionally, these early changes in neuronal morphology are potential contributors to the later functional changes observed in human iNeurons and *Df(h15q13)/+* cortical neurons. Taken together, these data reveal common deficits in mouse and human patient neuronal models of the 15q13.3 microdeletion, and a potential functional role for OTUD7A in disease pathogenesis.

### Elucidation of a neuron-specific OTUD7A protein-protein interaction network using proximity-labeling proteomics

Given the emerging clinical and genetic evidence supporting *OTUD7A* as a candidate driver of the 15q13.3 microdeletion and an independent NDD risk gene, we studied OTUD7A to understand neuron-specific signaling pathways. As little is known about OTUD7A, we used BioID2, a discovery-based proximity-labeling proteomics technique that has been widely used to identify protein-protein interactions and to map the spatial organization of cellular compartments^44, 46, 48, 73–76^. Proximity-labeling proteomics studies have typically used mammalian cell lines such as HEK 293 cells^47, 74, 76^, with only a few recent studies using this technique in neurons *in vitro* or *in vivo*^44, 46, 48, 73^. To identify OTUD7A, we optimized BioID2 in primary mouse cortical neurons using a lentiviral system (Fig. 3a). Neuron-specific expression of BioID2-fusion proteins was driven by a human Synapsin1 promoter, and neuron/glia co-cultures were used to promote synaptic maturation. TurboGFP was used as a reporter for transduction efficiency, followed by a self-cleaving P2A sequence to allow for bi-cistronic expression of TurboGFP and a gene-of-interest (GOI). This was followed by a flexible 13X GGGGS linker, to increase the biotinylation radius^51^, and a 3XFLAG tagged BioID2 sequence. We created lentiviral expression constructs for expression of OTUD7A-BioID2-3XFLAG, OTUD7A N492_K494del-BioID2-3XFLAG and OTUD7A L233F-BioID2-3XFLAG (**Supplementary Figure 6a**). Expression of BioID2 alone (BioID2-3XFLAG) provides a baseline list of background biotinylated proteins. To control for potential effects of overexpression, we created a negative control with an additional P2A sequence between the gene-of-interest and BioID2 (GOI-P2A-BioID2-3XFLAG); this was used for the majority of our BioID2 experiments (**Supplementary Figure 6a**). To ensure equal transduction efficiencies, lentiviruses were titered in primary cortical neurons prior to use. Mouse cortical neuron/ glia co-cultures transduced with BioID2-3XFLAG lentivirus displayed TurboGFP and FLAG expression in MAP2-positive neurons and not in GFAP-positive glial cells (**Supplementary Figure 6b**). All BioID2 fusion constructs were identified at their expected molecular weights in HEK293 FT cells and primary cortical neurons (**Supplementary Figure 6c,d**). Additionally, endogenous protein biotinylation was confirmed in transduced primary mouse cortical cultures (**Supplementary Figure 6e**).

**Figure 3.**
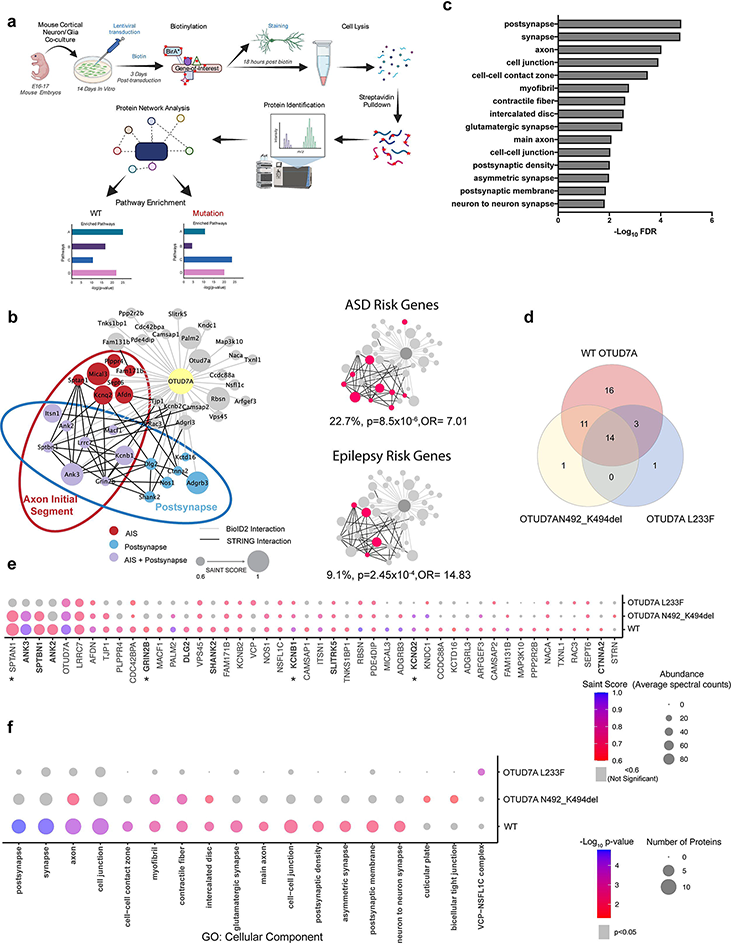
Neuron-specific BioID2 reveals an OTUD7A PPI network enriched for postsynaptic and axonal proteins which is impacted by patient mutations. **(a)** Workflow for lentiviral neuron-specific BioID2 experiment in mouse cortical neurons. **b)** Left: STRING analysis of functional interactions for WT OTUD7ABioID2 proteins with SAINT scores ≥ 0.6. Node size represents significance (SAINT SCORE). Node colours represent AIS (including those shared with AIS-BioID2 study by Hamdan et al., 2020^73^): red, postsynapse: blue, and proteins that are both AIS and postsynaptic: purple. Data shown are from 3 biological replicates. Right: Enrichment of ASD (SFARI Category 1/2/Syndromic) and epilepsy genes from Wang et al., 2017^63^. Fisher’s Exact Test. **(C)** Top 15 significant GO: Cellular component terms from functional enrichment analysis of the OTUD7A-BioID2 hits. p<0.05, Functional enrichment analysis was performed using gProfiler with Bonferroni correction for multiple testing. A custom background statistical domain scope was used (Sharma et al., 2015, mouse whole brain proteome)^62^ **(d)** Venn diagram of shared protein interactors between WT OTUD7A, OTUD7A N492_K494del and OTUD7A L233F. **(e)** Dotplot showing SAINT SCORE and average abundance of WT OTUD7A and OTUD7A patient mutation interactors. **(f)** Dotplot of enriched GO: Cellular component pathways in WT OTUD7A and patient mutation BioID2 hits.

To validate the neuronal BioID2 system, we used postsynaptic density 95 (PSD95), an excitatory synaptic scaffolding protein. Expression of PSD95-BioDI2-3XFlag in neurons displayed a punctate pattern along the dendrites that co-localized with biotinylated proteins identified via fluorescent streptavidin conjugates (**Supplementary Figure 6f,g).** We used the Significance Analysis of Interactome (SAINT)express tool to assign confidence scores to protein-protein interaction data^61, 77^ (**Supplementary Table 2**). SAINT scores range from 0 to 1, with a cutoff of 0.8 reliably predicting biologically relevant protein-protein interactions^77^. However, this cut-off was obtained using BioID data from cell lines utilizing very high numbers of cells with extremely homogeneous and reproducible samples, which would require a more stringent cut-off for false negatives. Due to the heterogeneous nature of mouse cortical cultures and the lower number of cells used in our pipeline, we used a SAINT score cut-off of 0.6 in order to avoid potential false negatives. We identified 147 PSD-95 interactors (**Supplementary Table 3**), 34 of which were shared with a PSD95 IP-MS study^78^ and 60 shared with an in vivo AAV PSD95-BioID2 study in mouse brain^46^ (**Supplementary Figure 6h**). Functional enrichment analysis revealed enrichment of PSD95 interactors in cellular compartments including the postsynaptic specialization, glutamatergic synapse and the postsynaptic membrane (**Supplementary Figure 6i; Supplementary Table 4**). These data demonstrate a strong degree of overlap with different PSD95 protein interaction studies; therefore, we moved forward with this approach to study OTUD7A protein interactors.

Transduced cortical neurons revealed a punctate localization pattern for OTUD7A-BioID2-3XFlag protein in the neurites (**Supplementary Figure 6f,g**), similar to our previous observation^30^, indicating that the BioID2 tag does not disrupt the localization of OTUD7A. Biotin and FLAG signals co-localized, further confirming proper proximity-dependent biotinylation (**Supplementary Figure 6f,g**). Following protein identification, SAINT analysis was performed using OTUD7A-P2A-BioID2 as the control. To create a comprehensive functional protein network map, we included all protein hits with a SAINT score of 0.6 and above (**Supplementary Table 3**). We identified a highly connected interactome of 44 proteins, which included postsynaptic density (PSD), cytoskeleton and axon initial segment (AIS) proteins, including 12 AIS proteins identified in a recent study that used BioID to map the axon initial segment (AIS)^73^ (Fig. 3b). Functional enrichment analysis revealed enrichment in cellular compartments including the postsynapse, synapse, axon, cell junction, cell-cell contact zone and postsynaptic density (Fig. 3c**; Supplementary Table 4**). Additionally, the top enriched molecular function GO terms for the OTUD7A PPI network were ion channel binding, spectrin binding, and structural constituent of the postsynaptic density, further implicating the cytoskeleton and postsynaptic density (**Supplementary Table 4**). These data are consistent with previous imaging studies showing that OTUD7A is localized to dendritic spines^30, 36^, but also identify the axon and AIS as novel cellular compartments occupied by OTUD7A. To determine whether the OTUD7A interactome includes known ASD-associated genes, we compared the OTUD7A PPI network with the gene list from the SFARI database of high-confidence ASD risk genes. We identified 10 high confidence category 1, 2 or syndromic SFARI genes in the list of 44 OTUD7A BioID2 hits (Fig. 3b; **Supplementary Table 5**) (22.7%; p=7.01×10^-5^, Fisher’s Exact Test; OR= 5.4), as well as 4 high-confidence epilepsy-associated genes^63^ (Fig 3b;**Supplementary Table 5**) (9.1%; p=2.45×10^-4^, Fisher’s Exact Test; OR=14.83), suggesting that OTUD7A may be a component of known ASD and epilepsy-associated pathways.

### Shared and Distinct Effects of patient mutations on functional OTUD7A PPI networks

To examine clinically relevant OTUD7A protein interactors, we compared the PPI networks of OTUD7A to the OTUD7A N492_K494del (ASD) and OTUD7A L233F (epileptic encephalopathy) mutations. The patient mutations showed a general decrease in the number of hits, with 26 in the OTUD7A N492_K494del list and 18 in the OTUD7A L233F list (**Supplementary Table 3**). There was a large degree of overlap between the mutation and WT OTUD7A PPI networks, with 14 proteins shared between all three conditions and only one protein unique to each of the mutations (Fig. 3d). To examine the impact of the mutations on individual protein interactions, we calculated the fold change of the abundance of each protein in the list for each mutation, compared to its abundance in the WT OTUD7A list (**Supplementary Table 6**). The OTUD7A N492_K494del list showed a greater than 30% decrease in the abundance of 10 proteins, with SHANK2 showing the greatest decrease (**Supplementary Table 6**, Fig. 3e). Protein abundances of the OTUD7A L233F interactors were decreased by as much as 80% for some proteins, indicating a severe impact on protein interaction (**Supplementary Table 6**, Fig. 3e). It is important to note that the abundance of OTUD7A itself was decreased by about 30%, which could be due to decreased expression of the OTUD7A L233F-BioID2 itself and/or reduced binding of the fusion protein to endogenous mouse OTUD7A (**Supplementary Table 6**). However, there was no difference in endogenous expression of OTUD7A in OTUD7A^L233F/L233F^ patient-derived iNeurons (Fig. 2l). In the OTUD7A L233F list, there were 26 proteins with a greater than 30% decrease in abundance, 13 of which are known or putative AIS proteins (**Supplementary Table 6**). This suggests that the OTUD7A L233F variant may significantly impact either localization of OTUD7A at the AIS or the direct binding of proteins at the AIS. Functional enrichment analysis of the OTUD7A N492_K494del list showed decreased or lost enrichment in most cellular compartments, most notably the postsynapse and synapse (Fig. 3f, **Supplementary Table 4**), whereas enrichment of ASD associated genes remained significant (19.2%; p=0.01, Fisher’s Exact test; O=4.34). Functional enrichment was lost for all cellular compartments in the OTUD7A L233F list, including the postsynapse, synapse and axon. Interestingly, enrichment in the axon was decreased but not lost in the OTUD7A N492_K494del list, indicating that these mutations may have shared and distinct changes in protein interactions and/or localization (Fig. 3f, **Supplementary Table 4**). Additionally, enrichment of ASD associated risk genes was lost in the OTUD7A L233F list (11.1%; p=0.24; Fisher’s Exact Test; OR=2.27), further suggesting a role for this mutation in NDD associated phenotypes.

### OTUD7A binds to Ankyrin-G and Ankyrin-B, and shares a common PPI network

For biological validation, we focused on Ankyrin-G (*ANK3*), as it showed high abundance, high significance (**Supplementary Table 3**) and its interaction with OTUD7A was affected by patient mutations (**Supplementary Table 6**, Fig. 3e). Additionally, genetic variants in *ANK3* have been associated with bipolar, schizophrenia, and ASD^79–82^. Ankyrin-G is a master regulator of the AIS, coordinating the precise organization of ion channels and structural proteins^83–89^. Ankyrin-G also regulates the structure and maturation of dendritic spines, through regulation in part by USP9X, a deubiquitinase (DUB) associated with neurodevelopmental disorders^54, 55^. We also validated Ankyrin-B, another Ankyrin family member that is strongly associated with epilepsy and ASD^90–93^. Like Ankyrin-G, Ankyrin-B acts as an adaptor and scaffold protein at the cell membrane, but is mainly localized to the dendrites and distal axon where it regulates axon growth, branching and trafficking^52, 94–97^. Additionally, the interaction between Ankyrin-B and OTUD7A was reduced by both patient mutations, suggesting that it may also contribute to 15q13.3 microdeletion neuronal phenotypes (Fig. 3a).

We examined the PPI network of Ankyrin-G by expressing an Ankyrin-G-BioID2 fusion protein in mouse cortical neurons. There are 3 major isoforms of Ankyrin-G in the brain, with the 270kDa and 480kDa isoforms being almost completely restricted to the AIS. The smallest isoform, at 190kDa, is the only isoform that has been identified in dendritic spines^53^. For this reason, and due to size restrictions of our lentiviral system, we used the 190kDa isoform. Following validation of the construct (**Supplementary Figure 7a-c),** we identified 12 Ankyrin-G-190 protein interactors in mouse cortical neurons, of which seven proteins were shared with the OTUD7A-BioID2 protein interactome (Fig. 4a,b), suggesting that OTUD7A and Ankyrin-G partially share a PPI network.

**Figure 4.**
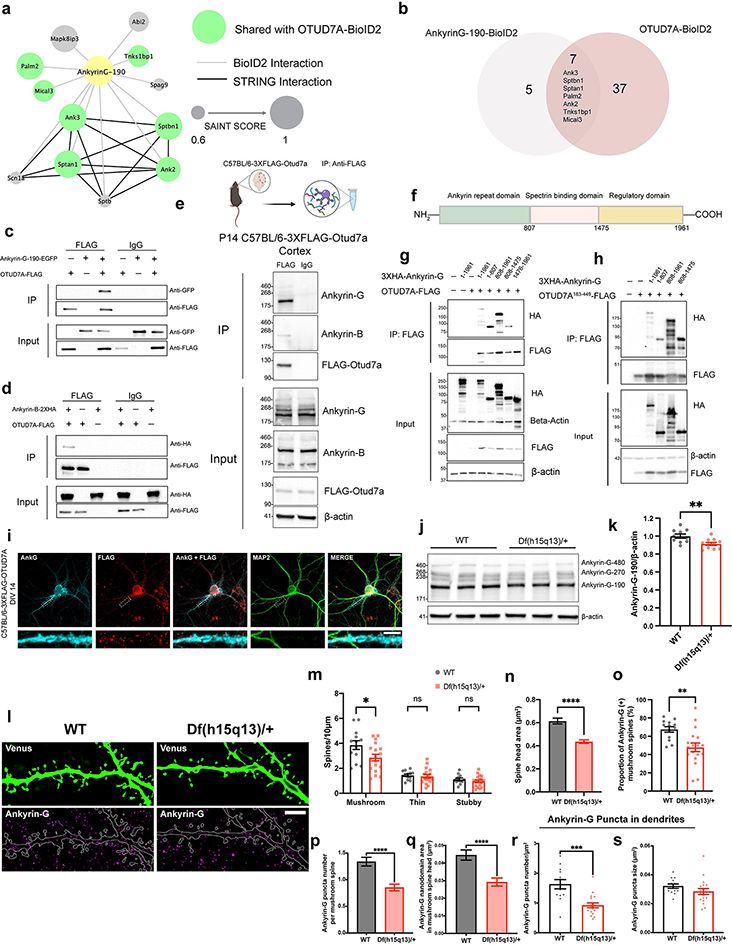
Ankyrin-G interacts with OTUD7A and its levels are decreased in *Df(h15q13)/+* mouse brain. **(a)** STRING analysis of functional network interactions for Ankyrin-G-190-BioID2 (from 3 biological replicates). Green nodes represent proteins shared with the OTUD7A-BioID2 list. **(b)** Venn diagram of shared protein interactors between Ankyrin-G-190 and OTUD7A. **(c)** Co-immunoprecipitation of OTUD7A-FLAG with Ankyrin-G-190-EGFP and **(d)** Ankyrin-B-2XHA from co-transfected HEK293 FT cells. **(e)** Co-immunoprecipitation of endogenous 3XFLAG-OTUD7A with Ankyrin-G and Ankyrin-B in P14 C57BL/6-3XFLAG-OTUD7A mouse cortex. **(f)** Schematic of mouse Ankyrin-G showing protein domains. **(g)** Domain mapping of Ankyrin-G-OTUD7A interaction from co-transfected HEK293FT cells expressing HA-Ankyrin-G domains and OTUD7A-FLAG or **(h)** the catalytic domain of OTUD7A (OTUD7A^183–449^-FLAG). **(i)** Representative image of DIV 14 C57BL/6J-3XFLAG-OTUD7A mouse cortical neurons stained for Ankyrin-G, FLAG, and MAP2. 63X objective; Top, scale bar = 20 μm; Bottom (Inset zoom), scale bar = 5 μm. **(j)** Representative western blot and **(k)** protein levels of Ankyrin-G-190 in P14 cortex from WT and *Df(h15q13)/+* mice; WT: n=10 cortices, *Df(h15q13)/+*: n=11 cortices; **p<0.01; student’s t-test, t=3.070, df=19. **(l)** Representative SIM images from DIV 17 WT and *Df(h15q13)/+* cortical neurons; Scale bar= 5 μm. **(m)** Spine morphology analysis. n= 13 WT and 18 *Df(h15q13)/+* cortical neurons (one dendrite per neuron). *p<0.05, Unpaired t-test (two tailed); Mushroom: t=2.301, df=29, p=0.0288; Thin: t=0.4023, df=29, p=0.6904; Stubby: t=0.9324, df=29, p= 0.3588. **(n)** Mushroom spine head. n= 322 spines WT, n= 350 spines Df(h15q13)/+. ****p<0.0001, Mann Whitney test two-tailed (approximate), U=40466; WT: median =0.4975, *Df(h15q13)/+*: median= 0.3680. **(o)** Proportion of Ankyrin-G (+) mushroom spines. **p<0.01, Student’s unpaired t-test; t=3.196, df= 29. **(p)** Ankyrin-G puncta number in mushroom spine heads. n= 322 spines WT, n= 350 spines Df(h15q13)/+. ****p<0.0001, Mann Whitney test two-tailed (approximate), U=44355; WT: median = 1, n=322; Df(h15q13)/+ median=0, n=350. **(q)** Ankyrin-G total nanodomain area in mushroom spines. Mann-Whitney test two-wailed (approximate), p=0.2204, U=18693; WT: median=0.046, n=244 spines; Df(h15q13)/+: median=0.052, n=165 spines. **(r)** Ankyrin-G puncta density in dendrites. ***p<0.001, Mann Whitney test two-tailed, p=0.004, U= 32; WT: median =1.548, n=13; *Df(h15q13)/+*: median + 0.8897, n=18. **(s)** Ankyrin-G puncta size in dendrites. Values are the average puncta size per dendrite. Mann Whitney test two-tailed (Exact), p=0.079, U=73; WT: median=0.03100, n=13; *Df(h15q13)/+*: median=0.02750,n=18.

We tested whether OTUD7A interacts with Ankyrin-G-190 and/or Ankyrin-B through co-immunoprecipitation of tagged proteins in HEK 293 cells. We found that both Ankyrin-G-190 and Ankyrin-B were co-immunoprecipitated with OTUD7A (Fig. 4c,d). To determine if these interactions exist in a more physiological context, we performed co-immunoprecipitation experiments in P20 C57BL/6-3XFlag-Otud7a mouse cortex. This newly created mouse line expresses endogenous N-terminal 3X flag tagged OTUD7A (**Supplementary Figure 7d**), with OTUD7A expression peaking at P21 in the cortex (**Supplementary Figure 7e,f**). Following pulldown of 3XFLAG-OTUD7A in cortical brain lysates, we found that all three neuronal isoforms of Ankyrin-G were co-immunoprecipitated with OTUD7A (with the 190kDa isoform showing the strongest signal) as well as the 220kDa isoform of Ankyrin-B (Fig. 4e). Staining of endogenous OTUD7A in DIV 14 3XFLAG-OTUD7A mouse cortical neurons revealed a punctate pattern of OTUD7A expression throughout MAP2-positive neurons, including the dendrites and axon. We also observed partial co-localization of OTUD7A and Ankyrin-G in the AIS of MAP2-positive neurons (Fig. 4i). We next identified the protein regions of Ankyrin-G that interact with OTUD7A through co-immunoprecipitation of overexpressed HA-tagged Ankyrin-G domains and OTUD7A-FLAG in HEK293 cells. We determined that OTUD7A binds to the N-terminal ANKRD, as well as the spectrin-binding domain, but not the C-terminal regulatory domain (Fig. 4f,g). Furthermore, we found that overexpression of the OTUD7A catalytic domain (OTUD7A^183–449^–FLAG) with HA-tagged Ankyrin-G domains was sufficient for binding to the ANKRD and spectrin-binding domains of Ankyrin-G (Fig. 4h). A previous study showed that both the ANKRD and spectrin-binding domain are ubiquitinated, whereas the regulatory domain is not^98^, suggesting that OTUD7A DUB function may modify these domains of Ankyrin-G. These data suggest that there is a strong interaction between OTUD7A and Ankyrin-G/ Ankyrin-B, and that OTUD7A may play a functional role at the ANKRD and/or spectrin-binding domain of Ankyrin-G.

### Ankyrin-G levels are altered in Df(h15q13)/+ mouse brain

To determine how OTUD7A may regulate Ankyrin-G in the mouse brain, we analyzed Ankyrin-G levels in WT and *Df(h15q13)/+* mouse cortex at P14. We observed a small (∼10%) but significant decrease in Ankyrin-G-190 levels in *Df(h15q13)/+* mouse cortex (Fig. 4j,k). We hypothesize that the regulation of Ankyrin-G by OTUD7A is neuron-specific, which may explain the small change in brain tissue. Therefore, we examined the subcellular localization of Ankyrin-G levels in mouse *Df(h15q13)/+* cortical neurons using structured illumination microscopy (SIM). A previous study using SIM imaging showed that Ankyrin-G forms condense clusters (nanodomains) that are organized peri-synaptically and in the spine head and neck^53^. We used the same SIM method to analyze the precise spatial organization of Ankyrin-G in the dendrites and dendritic spines of WT and *Df(h15q13)/+* mouse cortical neurons (Fig. 4l). Analysis of dendritic spines revealed that spine head size and mushroom spine density are decreased in *Df(h15q13)/+* neurons, corroborating our previous findings^30^ (Fig. 4m,n). We found that the proportion of mushroom spines expressing Ankyrin-G is decreased in *Df(h15q13)/+* neurons (Fig. 4o). Additionally, the number of Ankyrin-G puncta and the total Ankyrin-G nanodomain area per mushroom spine head was decreased in *Df(h15q13)/+* neurons (Fig. 4p,q). The density of Ankyrin-G puncta was also decreased in the dendrites of *Df(h15q13)/+* neurons whereas the size of Ankyrin-G puncta was unchanged (Fig. 4r,s). Given the interaction of OTUD7A with AIS-associated proteins, we analyzed Ankyrin-G intensity at the AIS and found a small but significant decrease in *Df(h15q13)/+* cortical neurons (**Supplementary Figure 8a,b**). Together these data indicate that Ankyrin-G levels are reduced in the dendrites, spines and AIS of *Df(h15q13)/+* mouse cortical neurons, which may underly the neuronal phenotypes in this mouse model.

### 15q13.3 microdeletion and OTUD7A^L233F/L233F^ patient iNeurons display alterations in Ankyrin-G protein levels, ubiquitination and stability

To determine if Ankyrin-G abnormalities extend to clinically relevant human models, we examined the subcellular localization of Ankyrin-G in iNeurons. SIM imaging revealed that Ankyrin-G intensity in the dendrites was decreased in iNeurons from the OTUD7A^L233F/L233F^ patient compared to the Family 1 control (Fig. 5a,b). Similar to the *Df(h15q13)/+* mouse model, both 15q13.3 microdeletion and OTUD7A^L233F/L233F^ iNeurons showed a decrease in the density of mushroom spines (Fig. 5a,c**; Supplementary Table 1**). However, mushroom spine head size was unchanged in both patient samples (Fig. 5a,d). Additionally, the number of Ankyrin-G puncta and the total Ankyrin-G nanodomain area in mushroom spine heads were decreased in both 15q13.3 microdeletion and OTUD7A^L233F/L233F^ iNeurons (Fig. 5a,e,f), recapitulating the SIM findings in *Df(h15q13)/+* mouse neurons. We examined Ankyrin-G staining intensity at the axon initial segment (AIS) and found a significant reduction in both 15q13.3 microdeletion and OTUD7A^L233F/L233F^ iNeurons (Fig. 5g,h).Together, these data suggest that Ankyrin-G levels are decreased in the dendritic spines and the AIS of patient neurons. Importantly, this phenotype is displayed by both 15q13.3 microdeletion and OTUD7A^L233F/L233F^ iNeurons, further implicating OTUD7A as a regulator of Ankyrin-G.

**Figure 5.**
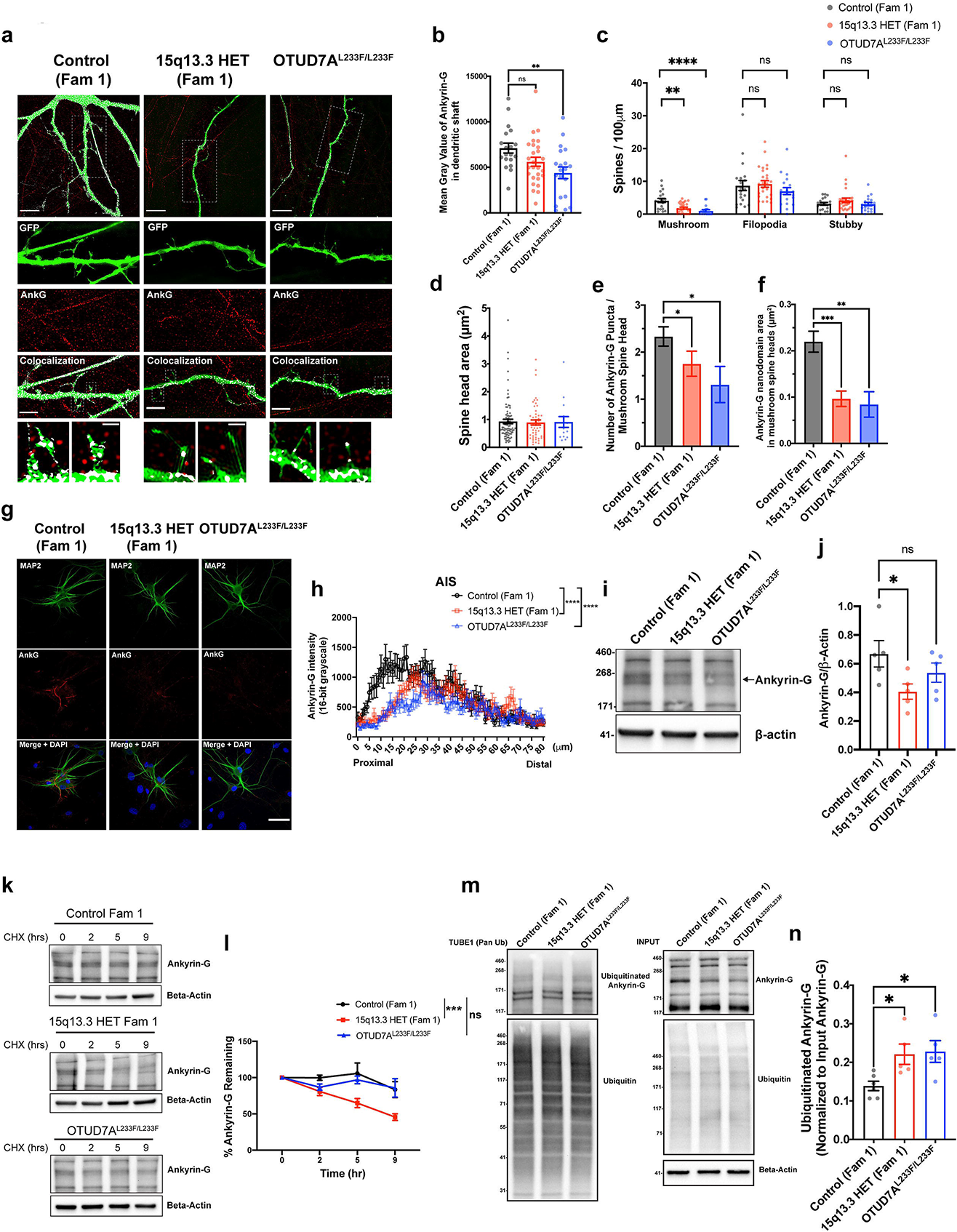
Ankyrin-G displays decreased levels and altered protein stability and ubiquitination in human 15q13.3 microdeletion and OTUD7A^L233F/L233F^ patient iNeurons. **(a)** SIM imaging of Venus-transfected PNI day 28 Family 1 and OTUD7A^L233F/L233F^ iNeurons stained for Ankyrin-G. Top: Scale bar = 10 µm, Middle: Scale bar= 5 µm, Bottom Scale bar = 1µm. **(b)** Ankyrin-G intensity in the dendrites (**p<0.01; One-Way ANOVA with Dunnett’s post-hoc test; F(2,63)=5.261, p=0.0077), **(c)** spine morphology (**p<0.01, ****p<0.0001; Mushroom: Kruskal-Wallis test with Dunn’s post-hoc test, p<0.0001 (approximate), Kruskal-Wallis statistic = 19.20; Filopodia: One-Way ANOVA with Dunnett’s post-hoc test, F (2, 63) = 0.9011, P=0.4113; Stubby: Kruskal-Wallis test with Dunn’s post-hoc test, p=0.5212 (approximate), Kruskal-Wallis statistic=1.303. Control (Fam 1) n= 20 dendrites, 15q13.3 HET (Fam 1) n=27 dendrites, OTUD7A^L233F/L233F^ n= 19 dendrites. **(d)** mushroom spine head area (One-Way ANOVA with Bonferroni’s post-hoc test; F(2,148)=0, 0=0.9658. **(e)** number of Ankyrin-G puncta in mushroom spine heads (Kruskal-Wallis test with Dunn’s post-hoc test; p=0.0167 (approximate Kruskal-Wallis statistic: 8.189) and **(f)** Ankyrin-G nanodomain area in mushroom spine heads. One-Way ANOVA with Dunnett’s post-hoc test; F(2,149)=10.06, p<0.0001); Control (Fam 1): n=84 spines, 15q13.3 HET (Fam 1) n= 53 spines, OTUD7A^L233F/L233F^ n=15 spines *p<0.05, **p<0.01, ***p<0.001. **(g)** Confocal images of Family 1 and OTUD7A^L233F/L233F^ PNI day 28 iNeurons stained for MAP2 and Ankyrin-G. Scale bar= 50 µm. **(h)** Ankyrin-G intensity (mean grey value) in the AIS. ****p<0.0001, Two-Way ANOVA with Dunnett’s post-hoc test; Interaction: F (160, 6196) = 1.891, P<0.0001; Distance from soma: F (80, 6196) = 8.481, P<0.0001; Genotype: F (2, 6196) = 69.18, P<0.0001. **(i)** Western blot and **(j)** analysis of Ankyrin-G in PNI day 7 human iNeurons from Family 1 and OTUD7A^L233F/L233F^. n= 5 separate Ngn2/Rtta transductions per line, *p<0.05, One Way ANOVA with Dunnett’s post-hoc test, F (2, 12) = 3.317, P=0.0713). **(k)** Western blot of time-course of Ankyrin-G levels after cycloheximide (20 µg/mL) treatment. **(l)** Kinetics of Ankyrin-G protein stability in Family 1 and OTUD7A^L233F/L233F^ induced neurons. n= 3 NGN2 transductions per condition; ***p<0.001; Simple Linear Regression followed by comparison of slopes by One-Way ANOVA with Dunnett’s post-hoc test; F (2, 30) = 10.99, P=0.0003. **(m)** TUBE pull-down from Family 1 and OTUD7A^L233F/L233F^ human iNeurons probed for Ankyrin-G and Ubiquitin. **(n)** Quantification of ubiquitinated Ankyrin-G from TUBE pulldown, normalized to the levels of Ankyrin-G in the input (whole lysate). n= 6 wells Control, 5 wells 15q13.3 microdeletion, 5 wells OTUD7A^L233F/L233F^; *p<0.05, One-Way ANOVA with Dunnett’s post-hoc test, F (2, 13) = 5.238, P=0.0215.

We further examined Ankyrin-G protein levels in patient iNeurons through western blotting. At PNI day 7, there was a significant reduction in the levels of a ∼200kDa isoform of human Ankyrin-G in 15q13.3 microdeletion iNeurons (Fig. 5i,j). This isoform has 95% protein sequence homology with mouse Ankyrin-G-190, which was significantly reduced in *Df(h15q13)/+* cortex (Fig. 4j,k). We hypothesized that Ankyrin-G stability may be affected in patient iNeurons, leading to the observed changes in Ankyrin-G protein levels. We treated PNI day 7 iNeurons with cycloheximide to block protein translation, allowing us to examine the degradation kinetics of Ankyrin-G. We found that there was an increase in the rate of Ankyrin-G-200 degradation in 15q13.3 microdeletion iNeurons compared to control iNeurons (Fig. 5k,l). This phenotype was not observed in OTUD7A^L233F/L233F^ iNeurons (Fig. 5k,l). In light of OTUD7A’s putative DUB function and our findings that the catalytic domain of OTUD7A binds to Ankyrin-G, we examined the ubiquitination status of Ankyrin-G in 15q13.3 microdeletion and OTUD7A^L233F/L233F^ iNeurons. Additionally, changes in degradation kinetics of proteins are often tightly linked to changes in ubiquitination status, as ubiquitinated proteins are often targeted for degradation by the proteasome^99^. Ubiquitinated proteins from PNI day 7 iNeuron lysates were pulled down using a tandem ubiquitin binding entity (TUBE) that binds to polyubiquitinated proteins. Analysis of Ankyrin-G levels in the polyubiquitinated pool of proteins revealed an increase in polyubiquitinated Ankyrin-G-200 in 15q13.3 microdeletion and OTUD7A^L233F/L233F^ iNeurons (Fig. 5m,n). Our data are consistent to some degree with a previous study that showed that OTUD7A^L233F/L233F^ patient fibroblasts displayed an increase in K48-polyubiquitinated proteins and proteasomal dysfunction^37^. We hypothesize that in OTUD7A^L233F/L233F^ iNeurons, although Ankyrin-G is highly poly-ubiquitinated and may be targeted for degradation, it may not be processed by the proteasome, which would explain the lack of change in protein stability. Additionally, these data are consistent with our finding that Ankyrin-G-200 protein levels are significantly decreased in 15q13.3 microdeletion iNeurons but not in OTUD7A^L233F/L233F^ iNeurons (Fig. 5i,j). Taken together, the data suggest that Ankyrin-G protein stability is decreased in 15q13.3 microdeletion iNeurons, which may be mediated by an increase in polyubiquitination of Ankyrin-G.

### 15q13.3 microdeletion and OTUD7A^L2^^33/L233F^ patient iNeurons display defects in intrinsic excitability and axon growth

Given the association of OTUD7A with axon-related proteins and the decrease in Ankyrin-G levels at the AIS, we were prompted to examine axon growth in patient iNeurons. We measured axon length in early patient iNeurons (PNI day 10), when individual axons can be distinguished using SMI-312 staining. We found that 15q13.3 microdeletion iNeurons showed a significant reduction in axon length, which was consistent in all three probands from three separate families (Fig. 6a-d**; Supplementary Table 1**). Our data suggest that neuronal connectivity defects associated with this CNV arise due to both axonal and dendritic growth abnormalities.

**Figure 6.**
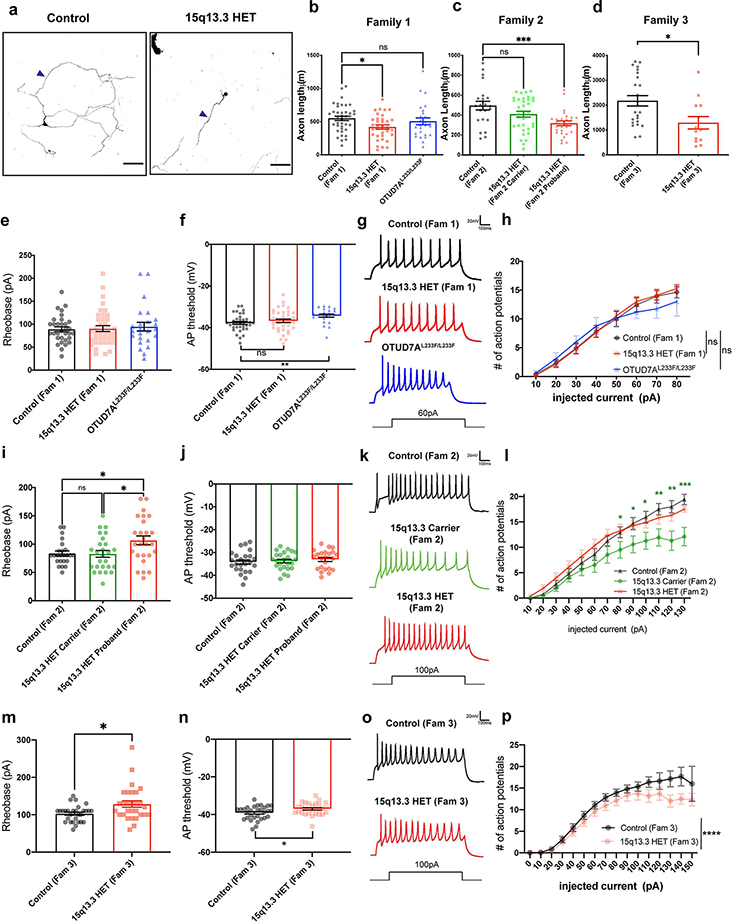
Human 15q13.3 microdeletion patient iNeurons display impairments in axon growth and intrinsic excitability. **(a)** Confocal images of PNI day 10 Venus-transfected Control and15q13.3 HET iNeurons. 20X Objective, tiled and stitched. Scale bar = 100 µm. **(b)** Quantification of axon length in Family 1 and OTUD7A^L233F/L233F^ iNeurons. n= 39 neurons Control Fam 1, n= 34 neurons 15q13.3 HET Fam 1, n=26 neurons OTUD7A^L233F/L233F^; *p<0.05, One-Way ANOVA with Dunnett’s post-hoc test, F (2, 96) = 3.306, P=0.0409. **(c)** Quantification of axon length in Family 2 iNeurons. n=23 neurons Control Fam 2, n= 34 neurons 15q13.3 HET Fam 2 carrier, and n= 28 neurons 15q13.3 HET proband Fam 2. ***p<0.001, One-Way ANOVA with Dunnett’s post-hoc test, F (2, 82) = 7.021, P=0.0015. **(d)** Quantification of axon length in Family 3 iNeurons. n=24 neurons Control Fam 3, n= 13 15q13.3 HET. *p<0.05, Unpaired t-test (two-tailed), t=2.614, df=35, p=0.0131. **(e-f)** Family 1 and OTUD7A^L233F/L233F^ iNeuron intrinsic membrane properties. **(e)** Rheobase (Kruskal-Wallis test with Dunn’s post-hoc test; Kruskal-Wallis statistic= 0.1493, p=0.9281 approximate; Control (Fam 1): n=32 neurons, 15q13.3 HET (Fam 1): n=38 neurons, OTUD7A^L233F/L233F^: n=24 neurons) and **(f)** action potential threshold (**p<0.01, One-Way ANOVA with Dunnett’s post-hoc test; F (2, 93) = 4.372, P=0.0153); Control (Fam 1): n=32 neurons, 15q13.3 HET (Fam 1): n=38 neurons, OTUD7A^L233F/L233F^: n=23 neurons. **(g)** Representative traces of action potentials evoked by 60 pA of injected current in PNI day 26-28 Family 1 and OTUD7A^L233F/L233F^ patient human iNeurons. **(h)** Repetitive firing properties of PNI day 26-28 Family 1 and OTUD7A^L233F/L233F^ patient human iNeurons. Two-Way ANOVA with Dunnett’s post-hoc test. Interaction: F (14, 325) = 0.4370, P=0.9620; Injected current: F (7, 325) = 74.84, P<0.0001; Genotype: F (2, 325) = 0.2437, P=0.7838. **(i-j)** Family 2 Intrinsic membrane properties. **(i)** Rheobase (*p<0.05, One-Way ANOVA with Tukey’s post-hoc test, F (2, 73) = 4.730, P=0.0117) and **(j)** action potential threshold (One-Way ANOVA with Tukey’s post-hoc test, F (2, 68) = 0.3923, P=0.6770). Control (Fam 2): n=24 neurons, 15q13.3 HET Carrier (Fam 2): n=26 neurons, 15q13.3 HET Proband (Fam 2): n=26 neurons. **(k)** Representative traces of action potentials evoked by 100 pA of injected current in PNI day 26-28 Family 2 human iNeurons. **(l)** Repetitive firing properties of PNI day 26-28 Family 2 human iNeurons. *p<0.05, **p<0.01, ***p<0.001, Two-Way ANOVA with Dunnett’s post-hoc test. Interaction: F (24, 666) = 1.253, P=0.1879; Injected current: F (12, 666) = 75.03, P<0.0001; Genotype: F(2, 666) = 32.16, P<0.0001. Control (Fam 2): n=24 neurons, 15q13.3 HET Carrier (Fam 2): n=25 neurons, 15q13.3 HET Proband (Fam 2): n=23 neurons. **(m-n)** Family 3 Intrinsic membrane properties. **(m)** Rheobase (*p<0.05, Mann-Whitney test two-tailed, Mann-Whitney U=271, p=0.0180; Control: 110.0 median, n=29 neurons, 15q13.3 HET: 120.0 median, n=29 neurons) and **(n)** action potential threshold (*p<0.05, unpaired t-test two-tailed; t=2.347, df=57, p= 0.0224; Control: n=30 neurons, 15q13.3 HET: n=29 neurons). **(o)** Representative traces of action potentials evoked by 100 pA of injected current in PNI day 26-28 Family 3 human iNeurons. **(p)** Repetitive firing properties of PNI day 26-28 Family 3 human iNeurons. Two-Way ANOVA with Sidak’s post-hoc test; Interaction: F (15, 603) = 0.7090, P=0.7766; Injected current: F (15, 603) = 78.61, P<0.0001; Genotype: F (1, 603) = 21.41, P<0.0001.

Neuronal excitability depends on proper AIS structure and composition, and loss of Ankyrin-G has been shown to result in deficiencies in the structure of the AIS and ability of neurons to initiate action potentials and fire repetitively^57, 84, 88, 89,100^; therefore, we examined whether there are changes to intrinsic membrane properties and excitability in patient iNeurons. We performed whole-cell patch clamp electrophysiology on PNI day 26-28 iNeurons (co-cultured on mouse glia) to identify electrophysiological abnormalities in individual neurons. We found that rheobase was increased in 15q13.3 microdeletion probands from Families 2 and 3 (Fig. 6i,m**; Supplementary Table 1**), and that the action potential threshold was increased in the15q13.3 microdeletion proband from Family 3 well as the OTUD7A^L233F/L233F^ proband (Fig. 6f,n**; Supplementary Table 1**) . There were no significant changes in other intrinsic membrane properties or action potential properties (Fig. 6e,j**; Supplementary Figure 9; Supplementary Table 1**). We examined repetitive firing using a step protocol and observed that, in the Family 3 15q13.3 microdeletion proband, there was a decrease in the number of action potentials fired (Fig. 6 p**; Supplementary Table 1**). Interestingly, we also observed this phenotype in the maternal carrier in Family 2 (Fig. 6l**; Supplementary Table 1**). The Family 1 and 2 probands showed no significant change in repetitive firing (Fig. 6h,l**; Supplementary Table 1**). The patch-clamp data suggest that patient iNeurons display a decrease in various intrinsic excitability parameters, which are consistent with the reduced AIS Ankyrin-G expression displayed by these neurons.

Taken together, our human iNeuron studies demonstrate that heterozygous deletion of the 15q13.3 region or a homozygous L233F point mutation in *OTUD7A* result in functional and morphological defects, along with reduced Ankyrin-G levels and stability, and increased ubiquitination. While the phenotypes are not expected to be identical between mouse and human iNeuron models, both models do show a level of convergence of molecular and cellular phenotypes, supporting the notion that disruption of the OTUD7A-Ankyrin-G interaction is a key pathogenic mechanism in the 15q13.3 microdeletion.

### Restoring OTUD7A or Ankyrin-G expression rescues morphological defects in mouse and human cellular models of the 15q13.3 microdeletion

To date, two candidate driver genes in the 15q13.3 microdeletion, *OTUD7A* and *CHRNA7*, have been the focus of attempts to rescue phenotypes in *Df(h15q13)/+* mice. We previously demonstrated that re-expression of OTUD7A *in vivo* rescues dendritic growth defects in mouse cortical neurons^30^, whereas another study stimulated CHRNA7 using an alpha7 positive allosteric modulator^101^. These studies suggest that loss of *CHRNA7* and *OTUD7A* may have additive or synergistic contributions to the 15q13.3 microdeletion. However, other cellular phenotypes have not been rescued in models of the 15q13.3 microdeletion, and neither have there been any rescues of phenotypes in patient neuronal models of the 15q13.3 microdeletion.

We first tested whether re-expression of OTUD7A in human 15q13.3 microdeletion iNeurons could rescue neuronal phenotypes. We co-transfected mCherry-tagged OTUD7A and VENUS in 15q13.3 microdeletion iNeurons from PNI day 21-28 and analyzed dendritic arborization (Fig. 7a). Sholl analysis revealed that re-expression of OTUD7A in15q13.3 microdeletion iNeurons increased dendrite arborization back to familial control levels, indicating that re-expression of OTUD7A is sufficient to rescue morphological defects in human 15q13.3 microdeletion glutamatergic neurons (Fig. 7b,c). Given our data supporting the importance of the OTUD7A/Ankyrin-G interaction in 15q13.3 microdeletion neuronal phenotypes, as well as the observed changes in Ankyrin-G levels and stability in 15q13.3 microdeletion models, we examined whether re-expression of OTUD7A in *Df(h15q13)/+* cortical neurons is sufficient to increase Ankyrin-G levels and rescue the protein homeostasis defects. We transduced *Df(h15q13)/+* cortical neurons with FLAG-tagged OTUD7A lentivirus, and probed lysates for Ankyrin-G (Fig. 7d). Expression of WT OTUD7A-3XFLAG significantly increased levels of Ankyrin-G-270 and Ankyrin-G-190 compared to *Df(h15q13)/+* neurons transduced with TurboGFP (Fig. 7e-g). To examine the effects of OTUD7A patient mutations on Ankyrin-G levels, we also examined Ankyrin-G levels in *Df(h15q13)/+* cortical neurons transduced with OTUD7A N492_K494del or OTUD7A L233F lentivirus. We first observed that levels of OTUD7A N492_K494del-3XFLAG was significantly higher than WT OTUD7A-3XFLAG (after normalizing for transduction efficiency), whereas levels of OTUD7A L233F-3XFLAG were not statistically different from WT OTUD7A-3XFLAG levels (Fig. 7h). Expression of OTUD7A N492_K494del or OTUD7A L233F was able to increase levels of Ankyrin-G-270 but did not increase levels of Ankyrin-G-190 (Fig. 7e-g), suggesting that patient mutations may specifically affect the ability of OTUD7A to regulate the 190kDa isoform of Ankyrin-G-190. These data suggest that OTUD7A can regulate Ankyrin-G levels in *Df(h15q13)/+* cortical neurons, and that a reduction of Ankyrin-G in the 15q13.3 microdeletion may underlie abnormal neuronal maturation.

**Figure 7.**
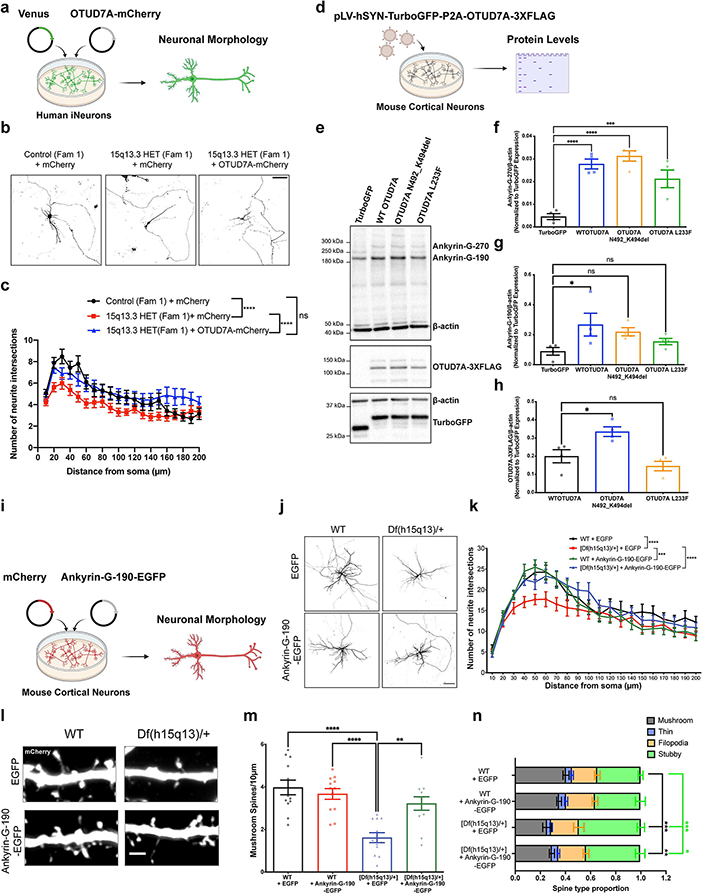
Ectopic expression of OTUD7A or Ankyrin-G rescues morphological impairments in human 15q13.3 microdeletion patient iNeurons or *Df(h15q13)/+* cortical neurons. **(a)** Family 1 control and 15q13.3 HET iNeurons were co-transfected with Venus and mCherry or Venus and OTUD7A-mCherry at PNI day 21 and fixed at for morphological analysis. **(b)** Representative confocal images of co-transfected Control and 15q13.3 HET Family 1 iNeurons, 20X objective, Scale bar= 100 µm. **(c)** Sholl analysis. n= 44 neurons Control + mCherry, n= 58 neurons 15q13.3 HET + mCherry, n= 45 neurons 15q13.3 HET + OTUD7A-mCherry; ****p<0.0001, Two-Way ANOVA with Tukey’s post-hoc test. Interaction: F (38, 2878) = 1.140, P=0.2565; Distance from soma: F (19, 2878) = 14.99, P<0.0001; Condition: F (2, 2878) = 43.39, P<0.0001. **(d)** *Df(h15q13)/+* cortical neurons were transduced with equal MOIs of TurboGFP, TurboGFP-P2A-WT OTUD7A-3XFLAG, TurboGFP-P2A-OTUD7A N492_K494-del-3XFLAG or TurboGFP-P2A-OTUD7A L233F-3XFLAG lentivirus at DIV 14 and lysed for western blotting at DIV 21. **(e)** Representative western blot and **(f)** levels of Ankyrin-G-270 (***p<0.001, ****p<0.0001, One-Way ANOVA with Dunnett’s post-hoc test, F (3, 12) = 20.96, P<0.0001) **(g)** Ankyrin-G-190 (*p<0.05, One-Way ANOVA with Dunnett’s post-hoc test, F (3, 12) = 3.169, P=0.0638 and **(h)** OTUD7A-FLAG (*p<0.05, One-Way ANOVA with Dunnett’s post-hoc test, F (2, 9) = 10.63, P=0.0043) in transduced *Df(h15q13)/+* primary cortical neurons; n= 4 transductions per genotype in 4 mouse cultures. **(i)** WT and *Df(h15q13)/+* mouse cortical neurons were co-transfected with mCherry and EGFP or mCherry and Ankyrin-G-190-EGFP at DIV 7 and fixed at DIV 14 for morphological analysis. **(j)** Representative images from DIV 14 WT and *Df(h15q13)/+* cortical neurons co-transfected with mCherry and EGFP or Ankyrin-G-190-EGFP; 20X Objective, scale bar 100μm. **(k)** Sholl analysis. n= 14 neurons WT + EGFP, n=15 neurons [Df(h15q13)/+] + EGFP, n= 13 neurons [Df(h15q13)/+] + Ankyrin-G-190-EGFP, n= 13 neurons WT + Ankyrin-G-190-EGFP, from 3 mouse cultures. ***p<0.001, ****p<0.000, 2-Way ANOVA with Tukey’s post-hoc test. Interaction: F (57, 1020) = 1.024, P=0.4272; DIV: F (19, 1020) = 38.23, P<0.0001; Genotype: F (3, 1020) = 16.88, P<0.0001. **(l)** Representative images of dendritic segments from co-transfected DIV 14 WT and *Df(h15q13)/+* cortical neurons; 63X Objective, scale bar 2 μm. **(m)** Mushroom spine density. **p<0.01, ****p<0.0001; One-Way ANOVA with Tukey’s post-hoc test; F (3, 44) = 13.04, P<0.0001. **(n)** Analysis of spine type proportions. n=12 neurons per condition from 3 mouse cultures. Two-Way ANOVA with Dunnett’s post-hoc test. Interaction: F (9, 176) = 4.682, P<0.0001; Condition: F (3, 176) = 0.001824, P=0.9999; Spine type: F (3, 176) = 146.5, P<0.0001.

To determine the direct role of Ankyrin-G-190 in morphological 15q13.3 microdeletion phenotypes, we transfected GFP-tagged Ankyrin-G-190 into WT and *Df(h15q13)/+* mouse cortical neurons and analyzed dendritic arborization and dendritic spines (Fig. 7i). Morphological analysis of DIV 14 *Df(h15q13)/+* neurons expressing GFP-tagged Ankyrin-G-190 showed a significant increase in dendritic arborization to a similar level as WT dendritic arborization (Fig. 7j,k). Expression of Ankryin-G-190 in a WT background did not affect dendritic tree complexity (Fig. 7j,k). Expression of Ankyrin-G-190 in *Df(h15q13)/+* cortical neurons also caused a significant increase in spine density, back to WT levels (**Supplementary Figure 10a**). More detailed analysis of spine morphology showed that mushroom spine density was also rescued by AnkryinG-190 expression (Fig. 7l,m). However, stubby spine density was also significantly increased in *Df(h15q13)/+* neurons (**Supplementary Figure 10b)** suggesting that Ankyrin-G-190 expression may be increasing spine formation without changing spine morphology. Corroborating this, aberrant spine type proportions in *Df(h15q13)/+* neurons were not rescued by Ankyrin-G-190 expression (Fig. 7n). Overexpression of Ankyrin-G-190 in a WT background did not significantly change any of the dendritic spine parameters analyzed. These data indicate that the effects of reduced Ankyrin-G levels may be important in early processes in neuronal development, including dendritic arborization and dendritic spine formation. The ability of Ankyrin-G to rescue neuronal deficits in 15q13.3 microdeletion neurons suggests that OTUD7A and Ankyrin-G may function in a common pathway to regulate phenotypes relevant to disease pathogenesis.

## DISCUSSION

The 15q13.3 microdeletion is a recurrent CNV associated with neuropsychiatric and neurodevelopmental disorders for which the pathogenic molecular mechanism(s) are unknown. Our study expands on previous work identifying *OTUD7A* as a novel driver gene in the 15q13.3 microdeletion^30^, which has since emerged as a novel and independent NDD risk gene^39–41^. However, the mechanism through which OTUD7A regulates disease-associated phenotypes is unknown^30, 36^. In this study, we used 15q13.3 microdeletion mouse neurons and human patient iPSC-derived induced neurons (iNeurons), along with OTUD7A^L233F/L233F^ iPSC-derived iNeurons, to reveal shared developmental impairments in cortical neuron activity and excitability, which were driven by reduced maturation of dendrites, dendritic spines and axons. Further, we developed a neuron-specific clinical mutation-based proximity-labeling proteomics assay and validation pipeline which revealed that OTUD7A interacts with a network of cytoskeletal, axonal and postsynaptic proteins. This network was enriched for ASD and epilepsy associated genes, including *Ank3* and *Ank2*. To validate our pipeline, we followed up on a top BioID2 hit, Ankyrin-G (*Ank3*), and found that OTUD7A interacts with Ankyrin-G and regulates its protein homeostasis. We found that Ankyrin-G had decreased protein stability, increased polyubiquitination, and reduced levels in dendritic spines and the AIS in 15q13.3 microdeletion iNeurons. Compellingly, many of these phenotypes were recapitulated in OTUD7A^L233F/L233F^ iNeurons, further highlighting the importance of OTUD7A in neural development and function. Given that we could rescue morphological impairments in 15q13.3 deletion neurons with OTUD7A or Ankyrin-G expression, our data reveal a key pathogenic impairment of OTUD7A-dependent regulation of Ankyrin-G in the 15q13.3 microdeletion.

Recent transcriptomic studies of the 15q13.3 microdeletion have implicated a wide array of potentially affected pathways^29, 31, 102^. However, changes in the transcriptome do not always align with changes to the proteome. Our proteomics approach identified an OTUD7A PPI network enriched in synaptic, cytoskeletal and axonal proteins, suggesting multiple developmental roles for OTUD7A in cortical neurons. This PPI network was also enriched for known NDD-associated proteins, including Ankyrin-G and Ankyrin-B, which are important regulators of the AIS, axon and dendritic spines. The interaction of OTUD7A with Ankyrin-G and Ankyrin-B were also affected by patient mutations, demonstrating the utility of using proximity-proteomics for mutational analysis, and the discovery of unexplored shared NDD disease mechanisms which are not captured using sequencing-based approaches. Our data reveal that OTUD7A interacts with Ankyrin-G and Ankyrin-B in the mouse cortex. Ankyrin-G is frequently referred to as the master regulator of the AIS, as depletion of Ankyrin-G results in drastic changes in AIS protein composition^84, 88^. We observed a decrease in the 190kDa isoform of Ankyrin-G in both mouse brain and human 15q13.3 microdeletion iNeurons, and SIM imaging revealed a decrease in Ankyrin-G nanodomains in dendritic spine heads. A previous study using SIM imaging found that the number of Ankyrin-G nanodomains positively correlates with spine head size and PSD size^53^, suggesting that it may modulate the PSD and contribute to the abnormal spine morphology observed in 15q13.3 microdeletion and OTUD7A^L233F/L233F^ neurons. We also found a decrease in Ankyrin-G expression at the AIS of human 15q13.3 microdeletion and OTUD7A^L233F/L233F^ iNeurons, which was accompanied by impairments in intrinsic excitability and axon length. Importantly, we observed both shared and distinct changes in electrophysiological properties between the three 15q13.3 microdeletion probands, highlighting the incomplete penetrance and heterogeneity displayed by this CNV. These data are consistent with the critical role of Ankyrin-G in AIS formation and maintenance as well as its role in axonal trafficking^85, 88, 89,103, 104^. Additionally, the AIS, dendrites and dendritic spine necks are enriched with the spectrin-actin cytoskeleton^105^ where OTUD7A and Ankyrin-G may help to stabilize these structures. Together, these data suggest that OTUD7A may drive developmental brain circuit deficits in the 15q13.3 microdeletion, which requires further examination in the future.

The OTUD7A protein is a member of the ovarian tumor (OTU) family of cysteine protease deubiquitinases (DUBs)^106, 107^. Substrate proteins are reversibly tagged with ubiquitin molecules through a series of enzymatic reactions; DUBs fall into this pathway by removing ubiquitin molecules from target proteins^108^. The most widely studied function of ubiquitination is protein turnover via the ubiquitin-proteasome system (UPS), which is a critical for learning and memory, and synaptic plasticity^108, 109^. Remarkably, OTUD7A and its family member OTUD7B are the only known DUBs showing linkage specificity to Lys 11 ubiquitin chains, although they can also target heterotypic chains containing Lys11/Ly48 or Lys11/Lys63 branched chains^107, 110, 111^. Lys11 chains typically do not target substrates to the UPS but are instead involved in signaling pathways^112^. However, heterotypic chains containing both Lys11/48 can target proteins for the UPS. Interestingly, a recent study showed that OTUD7A and OTUD7B interact with and modify Lys11/Lys63 heterotypic chains in order to regulate DNA damage^110^. This suggests that OTUD7A may regulate a wide range of cellular activities based on substrate linkage type. In a previous study, OTUD7A^L233F/L233F^ and OTUD7A KO HAP1 cells displayed proteasome dysfunction, identified by an increase in K48 polyubiquitinated proteins^37^. Additionally, pathogenic variants were identified in another OTU family DUB (OTUD6B) in 12 individuals with ID and seizures, which were also linked to reduced proteosome activity^113^. In the present study, we found that a polyubiquitinated pool of Ankyrin-G was increased in 15q13.3 microdeletion and OTUD7A^L233F/L233F^ iNeurons, indicating that proteasome dysfunction caused by this variant may also be present in neurons. However, we did not assay K48 polyubiquitin, so the changes that we observed may not be linked to proteasomal degradation. Additionally, rather than widespread proteasomal dysfunctional, the changes in ubiquitinated Ankyrin-G could also be a result of an impairment in direct OTUD7A DUB function. We found that Ankyrin-G protein stability was decreased in 15q13.3 microdeletion iNeurons, but not in OTUD7A^L233F/L233^ iNeurons. This result can potentially be explained by the proteasome dysfunction observed in OTUD7A^L233F/L233F^ cells^37^, such that polyubiquitinated Ankyrin-G may accumulate in the cells. Recent work has uncovered that another DUB, USP9X, regulates Ankyrin-G through its DUB activity^54, 55^; therefore, it is possible that OTUD7A may work together with USP9X to regulate Ankyrin-G levels and stability. The regulation of Ankyrin-G by OTUD7A and USP9X, and the link between Ankyrin-G and the spectrin-actin cytoskeleton indicates a potential critical convergence point during neuronal and circuit development. Human genetic sequencing studies have revealed that mutations in Ankyrin-G or spectrin proteins (SPTAN1, SPTBN4, SPTBN1) cause a range of NDDs^114–117^. Our findings of a shared OTUD7A-Ankryin PPI network including spectrin proteins suggests that a OTUD7A-Ankyrin-spectrin network could be important for the integrity and growth of developing axons and dendrites. Overall, our results suggest that OTUD7A may regulate Ankyrin-G proteostasis, and more broadly adds to the growing importance of the ubiquitin pathway, and DUBs in particular, in brain development and neurodevelopmental disorders^91, 108, 109, 118–124^.

Finally, we found that re-expression of either OTUD7A or Ankyrin-G in 15q13.3 microdeletion iNeurons or mouse neurons, respectively, was sufficient to rescue neuronal morphological phenotypes. These results indicate a central role for a conserved OTUD7A-Ankyrin-G interaction in neuronal dysfunction in the 15q13.3 microdeletion, providing molecular insight into disease processes.

Our study focused on *OTUD7A*, given emerging evidence that it is an independent NDD risk gene^39–41^, and our previous work showing evidence of its role as a driver gene in the 15q13.3 microdeletion^30^. However, it is likely that other genes in the 15q13.3 CNV also contribute. For example, *CHRNA7* was the first gene suspected to contribute to the 15q13.3 deletion^19, 125^, but follow up studies are conflicting with regards to the importance of *CHRNA7*^34, 126^. While we know that cortical excitatory neurons are dysfunctional in the 15q13.3 deletion, both *OTUD7A* and *CHRNA7* are also expressed in inhibitory neurons^127, 128^, and inhibitory neuron defects have been identified in 15q13.3 microdeletion mouse models^26, 29, 129^. Given the Ankyrin-G deficits at the AIS and the reduced intrinsic excitability in 15q13.3 microdeletion iNeurons, we speculate that inhibitory synapse formation onto the AIS of excitatory neurons could be a critical site of dysfunction. Other potential contributing genes with evidence for roles in neurodevelopmental phenotypes include *FAN1*, *KLF13*, and *TRPM1*^130–132^. It is also possible that these potential candidate genes may be acting in different cell populations and/or at different time points during brain development. Equally plausible is that different driver genes have prominent roles in different individuals with the deletion, or modifier mutations outside of the region as has been found for the 16p11.2 CNV^133, 134^.

Large recurrent CNVs such as the 15q13.3 microdeletion are complex and likely have multiple mechanisms of disease; however, the focus on driver genes such as *OTUD7A* is a critical starting point to dissect relevant signaling pathways for therapeutic intervention. While used in the context of a single CNV, our proteomics approach has broad applicability since there are multiple driver genes for other NDD-related CNVs. Pairing this analysis with independent disease-linked variants in candidate genes is a path forward to identify clinically relevant pathways. Further, the identification of critical driver genes in CNVs synergizes with emerging gene-therapy technologies^135^. Our study reveals that using proximity-labeling proteomics in a physiologically relevant cell type combined with multiple validation models and techniques can help to unravel complex biological pathways associated with recurrent CNVs.

## METHODS

### Animals

Df(h15q13)/+ mice were generated by Taconic Artemis as described previously^24^. Animals were bred, genotyped, housed and approved for use at the Central Animal Facility at McMaster University and the Animal Research Ethics Board (AREB), and by the Animal Resource Center at the University Health Network in Toronto, Canada. Genotypes were identified during breeding by PCR of ear notches, and two WT females were bred with 1 *Df(h15q13)/+* male per breeding cage. The use of only WT females for breeding was performed to minimize effects of potential differences in the embryonic environment and/or mothering of the *Df(h15q13)/+* females compared to WT females. To obtain cortical cultures, WT females were timed-bred with *Df(h15q13)/+* males and males were removed when a plug was observed, indicating copulation. At E16, mothers were sacrificed, and litters were collected. Animals of appropriate genotype were included, and any animals with unclear genotypes were excluded from experiments. The mouse line C57BL/6J-Otud7A<em1Tcp> (3XFLAG-Otud7a) was made at The Centre for Phenogenomics by electroporating Cas9 ribonucleoprotein complexes with a guide RNA with the spacer sequence GCTAGAGACCATCCATCTGC and a single-strand oligonucleotide encoding a 3XFLAG tag and GGSG flexible linker inserted immediately after the start codon. Also identified was an upstream intronic variant 1-bp delG Chr7:63650782 (GRCm38). For BioID2 studies, timed-pregnant CD1 mice were ordered from Charles River and were euthanized for embryo collection at E16.

### Cell Culture

Primary cortical neurons were cultured as follows. Cortices were dissected out of WT and Df(h15q13)/+ mouse embryonic brains at E16. The last 1-2mm of their tails were used for genotyping. Dissociation of cortices was aided by incubation in 0.3 mg/mL Papain (Worthington Biochemical)/400 U/mL DNase I (Invitrogen) in Hanks Buffered Saline Solution (HBSS) for 20 min at 37 °C, followed by light trituration. Cells were seeded onto 0.1 mg/mL poly-D-lysine (BD Sciences)/3.3 mg/mL Laminin (Sigma)-coated coverslips in plating media containing Neurobasal medium, 10% fetal bovine serum, 1% penicillin/streptomycin, and 2 mM GIBCO Glutamax supplement. After 1.5 hr, media was changed to serum-free feeding media containing Neurobasal medium, 2% B27 supplement, 1% penicillin/streptomycin, and 2 mM L-glutamine. For certain immunocytochemistry experiments, cultures were treated with 1 mM Cytosine b-D-arabinofuranoside hydrochloride (Ara-C) (Sigma) at DIV 4 to inhibit glial cell proliferation. Cultures were maintained at 37 °C, 5% CO2. All media components were from GIBCO unless otherwise specified.

HEK293 FT cells were maintained in DMEM with 4.5g/L glucose, 10% Fetal bovine serum, 2mM GIBCO Glutamax supplement, 1mM Sodium pyruvate and 1X MEM Non-essential amino acids. Lenti-X 293T cells were maintained in DMEM with 4.5g/L glucose, 10% Fetal bovine serum, 4mM GIBCO Glutamax supplement and 1mM Sodium Pyruvate.

### Antibodies and Constructs

The following antibodies were used in this study. Mouse anti-FLAG (Sigma F1804; Western Blot 1:1000, IF 1:1000; IP 1-4 µg), Mouse anti-HA (Santa Cruz Biotechnology F-7; Western Blot 1:500), Rabbit anti-Ankyrin-G (Synaptic Systems 386 003; Western blot 1:4000, IF 1:1000), Mouse anti-Ankyrin-G (Neuromab N106/36; IF 1:200) Mouse anti-ß-actin (Sigma A5316; Western blot 1:5000), Rabbit anti-ß-actin (Cell Signaling Technologies #4907; Western blot 1:1000), Rabbit anti-GFP (Santa Cruz Biotechnology sc-8334; Western blot 1:1000), Rabbit anti-TurboGFP (Thermo Fisher Scientific PA5-22688; Western blot 1:1000; IF 1:1000), Rabbit anti-mCherry (Abcam ab167453; IF 1:500), Chicken anti-GFP (Aves Labs AB_2307313; IF 1:1000), Rabbit anti-GFP (Invitrogen A-11122; Western Blot 1:1000), Rabbit anti-OTUD7A (MilliporeSigma HPA044554; Western Blot 1:500), Mouse anti-SMI-132 (Biolegend 837904; IF, 1:1000), Chicken anti-MAP2 (Cedarlane CLN 182; IF: 1:1000), Chicken anti-MAP2 (AVES labs MAP; IF 1:1000), Rabbit anti-Ubiquitin [EPR8830] (Abcam ab134953). Fluorophore-conjugated secondary antibodies were raised in donkey and used at a concentration of 1:1000. Alexa Fluor 647-Streptavidin (Jackson Immunoresearch; IF 1:1000) was used to detect biotin in IF experiments. Pierce™ High Sensitivity Streptavidin-HRP (Thermo Fisher scientific; western blot 1:30 000) was used to detect biotin in western blots. Tube 1 Magnetic Beads (Life Sensors UM401M) were used for ubiquitin pull-down experiments.

To create the BioID2 fusion constructs, we obtained an expression plasmid containing a C-terminal 3Xflag tagged BioID2 sequence with a 198bp (13X “GGGGS” repeat) linker sequence upstream of BioID2 (Genscript). For lentiviral expression, 13X linker-BioID2-3XFLAG was amplified and cloned into the lentiviral backbone pLV-hSYN-RFP^59^ (Addgene Plasmid #22909) using InFusion cloning. For ease of visualization and to create a bicistronic construct, TurboGFP-P2A was amplified from pCW57-GFP-2A-MCS (Addgene plasmid #71783) and cloned into the pLV-hSYN-RFP backbone between the BamHI and PmeI restriction sites, replacing RFP. InFusion cloning was used to insert individual transgenes between the P2A and the 13Xlinker-BioID2 sequences.

The pcDNA-OTUD7AL233F-FLAG construct was made by incorporating a C>T mutation at bp 697 of the human OTUD7DA cDNA sequence into the PCR primers prior to amplification. The PCR product was subcloned into the pcDNA3.3 backbone using the InFusion cloning system between the EcoRI and MluI restriction sites. OTUD7A^183–449^-FLAG was created through PCR amplification of the OTUD7A catalytic domain (bp547-1347 of human OTUD7A cDNA sequence) followed by subcloning into the pcDNA3.3 backbone using the InFusion cloning system between the EcoRI and MluI restriction sites. mCherry-tagged constructs were created by amplification of WT OTUD7A, OTUD7AN492_K494del or OTUD7A L233F from the pcDNA3.3 backbone and amplification of mCherry from Lenti U6-sgRNA EF1a-mCherry (Dr. Jeremy Day, UAB, Alabama), followed by InFusion of PCR products into the pcDNA3.3 backbone between the EcoRI and KpnI restriction sites. The pcDNA3.3 mCherry construct was created similarly by amplification of mCherry from Lenti U6-sgRNA EF1a-mCherry (Dr. Jeremy Day, UAB, Alabama), followed by InFusion of PCR products into the pcDNA3.3 backbone between the EcoRI and KpnI restriction sites.

Ankyrin-G-190-GFP (plasmid #31059) and Ankyrin-B-2XHA (plasmid #31057) were bought from Addgene. The 3XHA-Ankyrin-G domain constructs were gifts from Dr. Peter Penzes (Northwestern University). pCAGIG-Venus was provided by Dr. Zhigang Xie (Boston University, MA).

### iPSC Reprogramming

All pluripotent stem cell work was approved by the Canadian Institutes of Health Research Stem Cell Oversight Committee. Blood was collected from individuals with the approval from SickKids Research Ethics Board after informed consent was obtained, REB approval file 1000050639. This study was also approved by the Hamilton Integrated Research Ethics Board, REB approval file #2707. CD34+ blood cells were verified using flow cytometry and collected for iPSC reprogramming. iPSCs from Family 1 and 2 were reprogrammed by CCRM (MaRS centre, Toronto, ON) and iPSCs from family 3 were generated in-house. iPSCs were generated by Sendai virus reprogramming and clonal expansion using the CytoTune – iPSC 2.0 kit (ThermoFisher) to deliver the reprogramming factors. Once colonies were large enough (approximately 15-17 days post Sendai transduction), each colony was transferred to 1 well of a 12-well plate coated with irradiated MEFs and plated in iPSC media (DMEM/F12 supplemented with 10% KO serum, 1x non-essential amino acids, 1x GlutaMAX, 1mM β mercaptoethanol, and 16 ng/mL basic fibroblast growth factor (bFGF)). Once iPSCs were expanded and established, they were transferred to Matrigel-coated plates and grown in mTeSR1 (STEMCELL Technologies). ReLeSR (STEMCELL Technologies) was used for subsequent passaging. iPSC lines were validated through flow cytometry of pluripotency markers and G-banding analysis to confirm a normal karyotype.

### iPSC NGN2/Rtta infection

Singularized iPSCs were plated on Matrigel (Corning) coated 6-well plates at a density of 2.50 x 10^5^ cells per well, in 2mL of mTeSR (STEMCELL Technologies) containing 10 µM Y-27632 (STEMCELL Technologies). The following day, they were changed into 2mL of fresh mTeSR containing 10 µM Y-27632 and 1µg/mL polybrene, as well as Ngn2 and rtTA lentiviruses which had been titered and an MOI of 1 was used for all transductions. The virus containing media was replaced with fresh mTeSR at 24 hours post infection. The cells were allowed to recover to approximately 70-80% confluency before being passaged into 6-well Matrigel coated plates. At their next 70-80% confluency, cells were either frozen (in media containing 50% knockout serum (Invitrogen), 40% mTeSR, and 10% DMSO), or singularized for induction into iNeurons. All infected iPSCS were used within 5 passages of infection to maintain induction efficiency.

### iNeuron induction protocol

At Day -1 of induction, singularized Ngn2/rtTA infected iPSCs were plated at a density of 5.0 x 10^5^ cells per well onto Matrigel coated 6 well plates, in 2mL of mTeSR containing 10 µM Y-27632. At Day 0 of induction, cells were changed into fresh mTeSR containing [LOOK UP] µg/mL of doxycycline hyclate. On Days 1 and 2 of induction, cells received fresh iNPC media (DMEM/F12 containing 1% N-2 supplement, 1% Penicillin/Streptomycin, 1% NEAA, 1% Sodium Pyruvate and 1% GlutaMAX) containing 1µg/mL of doxycycline hyclate and 1-2 µg/mL puromycin. On Day 3, cells were changed into fresh iNI media (Neurobasal with SM1 supplement, 1% Penicillin/Streptomycin, and 1% GlutaMAX) containing BDNF (10 ng/µL; Peprotech), GDNF (10 ng/µL; Peprotech), and laminin (1µg/mL). On Day 4, cells were replated onto polyornithine/laminin coated 12mm coverslips, or polyethylenimine (PEI) coated 48-well multi-electrode array plates. On Day 5, mouse glia were added to the wells. All wells received half media changes of iNI with BDNF, GDNF and laminin for the duration of the experiment, with the inclusion of 2.5% FBS (v/v) starting at day 10. iNeurons that were used for RNA/protein extraction were maintained in their induction wells until Day 7 with no glia and received a fresh media change on Day 5.

### iNeuron RT-qPCR

iNeurons were plated without glia on 6-well plates at a density of 5 x 10^5^ cells per well. Total RNA was extracted at PNI day 7 using a commercial kit (Norgen, Cat. #17200), and 1µg of RNA was used for cDNA synthesis using the qScript cDNA synthesis kit according to manufacturer instructions (Quanta Biosciences). Primers were designed using the Universal Probe Library (UPL) Probefinder software for human (Roche, version 2.53) or using previously published sequences and adjusted to be intron-spanning. Quantitative PCR was performed using SYBR Green super mix (FroggoBio) and the QuantStudio 3 thermocycler (Applied Biosystems). Data were analyzed using the Thermo Cloud^TM^ to generate relative expression normalized to the housekeeper *EIF3L*.

### Multielectrode Arrays

48-well Cytoview MEA plates (Axion Biosystems, M768-tMEA-48B) were coated with 0.1% Polyethylenimine (PEI) 24 hours prior to plating. Primary mouse cortical cultures were dissected at E16 and plated onto PEI-coated 48-well Cytoview MEA plates at a density of 3 x 10^4^ cells/ well in plating media containing Neurobasal medium, 10% fetal bovine serum, 1% penicillin/streptomycin, and 2 mM GIBCO Glutamax supplement. After 1.5 hr, media was changed to serum-free feeding media containing Neurobasal medium, 2% B27 supplement, 1% penicillin/streptomycin, and 2 mM L-glutamine. Half of the culture media was replaced every 2 days.

For human iNeurons, 40,000 cells were plated on each well of a 48-well MEA plates on day 4 PNI and allowed to attach for 1.5 hours in a 37 degree Celsius incubator. On PNI day 5, 20,000 CD1 mouse glia were added to each well. The plates received half media changes every other day (iNI+BDNF+GDNF+laminin, with 2.5% FBS added after PNI Day 10). MEA recordings were taken approximately twice per week using the Axion Maestro Pro. Days *in vitro* (DIV) for iNeuron MEA recordings refers to the number of days after plating iNeurons onto the MEA plates. The day of plating (DIV 0) refers to PNI Day 4.

To record extracellular spontaneous activity, MEA plates were placed in the MaestroPro MEA system (Axion Biosystems) at 37 °C for 5 minutes to acclimate, followed by a 10-minute recording period. Spike data were analyzed with the AxIS Navigator software (Axion Biosystems) at a sampling rate of 12.5 kHz with a 4kHz Kaiser Window low pass filter and a 200Hz IIR High Pass filter. Analyzed data were exported to CSV files and statistical analysis was performed in GraphPad Prism 9 software. Wells that had zero active electrodes throughout the duration of the experiment were excluded for statistical analysis. Raster plots were generated with the Neural Metric Tool (Axion Biosystems).

### Electrophysiology

On Day 4 post induction, iNeurons were added to 24 well plates (Corning) at 100,000 cells/well on polyornithine/laminin coated 12mm coverslips, in iNI media containing BDNF, GDNF and laminin. On PNI day 5, 50,000 previously cultured CD1 mouse glia were plated onto these coverslips. The cells were maintained until for 4 weeks post-induction, receiving half media changes every other day. At PNI day 10, 2.5% FBS was added to the media and maintained for the duration of the experiment. Whole cell recordings (Olympus BX51 WI) were performed at room temperature using an Axoclamp 700B Amplifier (Molecular Devices). Trace recordings were performed using Clampex 10.7 and analyzed in Clampfit 10.7. Borosilicate glass pipettes (WPI; 1B150F-4) were used to prepare patch electrodes (P-97 or P1000; Sutter Instruments) and used for recordings with an intracellular solution containing (in mM): 123 K-gluconate, 10 KCl, 10 HEPES, 1 EGTA, 2 MgCl_2_, 0.1 CaCl_2_, 1 MgATP, 0.2 Na_4_GTP (pH 7.2 by KOH), with or without 0.06% (m/v) sulpharhodamine B to aid in visual identification of neuron morphology post-recording. Standard HEPES aCSF was used for all recordings, containing (in mM): 140 NaCl, 2.5 KCl, 1.25 NaH_2_PO_4_, 1 MgCl_2_, 10 glucose, 2CaCl_2_ (pH 7.4 by NaOH). Recordings were sampled at 10-20kHz and low pass filtered at 1 kHz. The membrane potential was clamped at −70mV adjusted for a junction potential of −10mV, and action potentials were elicited with step currents of 10 pA (beginning at −40pA). Recordings were excluded if the series resistance exceeded 25 MΩ

### Transfection

Primary mouse cortical neurons were transfected at DIV 7 using Lipofectamine LTX with Plus reagent (Invitrogen) according to the manufacturer’s instructions. Approximately 9×10^5 cells per well were transfected with 1µg DNA with 2 µL Lipofectamine LTX and 1µL Plus reagent. HEK293FT cells were grown under standard cell culture conditions and transfected with plasmids using Lipofectamine 2000 according to the manufacturer’s protocol (Invitrogen) at approximately 80-90% confluency. HEK293FT cells were used for ease of plasmid expression and have not been tested for Mycoplasma contamination. For transfection of iNeurons, neurons were plated at a density of 1.3 x 10^4^ or 5.3 x 10^4^ per cm^2^ with 2.6 x 10^4^ mouse glia per cm^2^, onto 12mm round coverslips coated with Poly-ornithine and laminin. For axon analyses, 350ng of pCAGIG-VENUS was used to transfect the neurons two days after adding mouse glia (PNI day 7) using Lipofectamine 2000 (ThermoFisher). A full media change using conditioned media was performed 5-6 hours post-transfection. Neurons were fixed with 4% paraformaldehyde 72 hours after transfection (PNI day 10) for downstream immunofluorescence. For sholl analysis, neurons were transfected with 125ng pCAGIG-VENUS and 375ng pcDNA-mCherry or pcDNA-OTUD7A-mCherry on PNI day 21 using Lipofectamine 2000. Cells were fixed with 4% PFA on PNI day 28 (7 days post-transfection).

### Lentivirus Production

Lenti-X 293T cells were transfected with transfer plasmid containing overexpression construct, along with the packaging plasmid psPAX2 (Addgene Plasmid #12260) and the VSV-G envelope plasmid PMD2.G (Addgene Plasmid #12259). 48 hours after transfection, viral supernatant was collected and ultracentrifuged at 25 000 RPM for 2 hours at 4°C. To titer the lentivirus, primary mouse cortical neurons were transduced at DIV 3 at three different concentrations, followed by flow cytometry at DIV 5 to calculate the percentage of GFP+ cells. Titers were calculated using the following equation: Fraction GFP+ × dilution × cell# / volume (mL) = transducing units (TU) / mL. Functional titers were calculated as the average titer obtained from the three dilutions and ranged from 10^7 – 10^8 TU/mL.

### *In vitro* Biotinylation

Cultured mouse primary cortical neurons (approximately 7.2 x10^6 neurons per virus) were transduced with lentiviral BioID2 constructs at a MOI of 0.7. Neurons were transduced at DIV 14 and biotin was added at DIV 17 at a final concentration of 50µM. Cells were lysed in RIPA buffer, sonicated, and biotinylated proteins were pulled down with streptavidin-sepharose beads (GE ref# 17-5113-01). Bead-protein conjugates were resuspended in ammonium bicarbonate and on-bead trypsin digestion was performed overnight at 37 °C. Beads were washed with ammonium bicarbonate and supernatants containing digested peptides were speed vac dried before preparation for LC-MS.

### Liquid Chromatography and Tandem Mass Spectrometry (LC-MS/MS) for BioID2

BioID2 samples were resuspended with 20 µl 0.1% formic acid, 1 ml out of 20 ml was injected for LC-MS/MS analysis. Liquid chromatography was conducted using a home-made trap-column (5 cm x 200 mm inner diameter) and a home-made analytical column (50 cm x 50 mm inner diameter) packed with Reprosil-Pur 120 C18-AQ 5 µm particles (Dr. Maisch), running a 3hour reversed-phase gradient at 70nl/min on a Thermo Fisher Ultimate 3000 RSLCNano UPLC system coupled to a Thermo QExactive HF quadrupole-Orbitrap mass spectrometer. A parent ion scan was performed using a resolving power of 120,000 and then up to the 30 most intense peaks were selected for MS/MS (minimum ion counts of 1000 for activation), using higher energy collision induced dissociation (HCD) fragmentation. Dynamic exclusion was activated such that MS/MS of the same m/z (within a range of 10ppm; exclusion list size=500) detected twice within 5s were excluded from analysis for 50s.

### Mass Spectrometric Data Analysis

Mass spectrometric raw files from the Thermo QExactive HF quadrupole-Orbitrap were searched using Proteome Discoverer, against the UniProt Mouse database (Version 2017-06-07), in addition to a list of common contaminants maintained by MaxQuant^60^. The database parameters were set to search for tryptic cleavages, allowing up to 2 missed cleavage site per peptide, with a parent MS tolerance of 10 ppm for precursors with charges of 2+ to 4+ and a fragment ion tolerance of ±0.02 amu. Variable modifications were selected for oxidized methionine. The results from each search were statistically validated within Proteome Discoverer, with 1 unique peptide and an FDR cutoff at 0.01 required for protein identification. SAINTexpress was used to calculate the probability of each potential proximal– protein from background (control BioID2) using default parameters^61^.

### PPI Network Analysis

Proteins with a SAINT score of greater than or equal to 0.6 were included in downstream analyses. Protein networks were constructed in Cytoscape (v3.8.2) using the STRING protein interaction database for mus musculus. Dotplots were created in RStudio. Functional enrichment tests were performed on gProfiler using the Bonferroni correction for multiple comparisons. A mouse brain proteomics reference list^62^ was used as a custom statistical domain scope. GO terms with FDR <0.05 were considered significant. For enrichment of NDD risk genes, we compared our BioID2 lists with a list of 657 category 1,2 and syndromic ASD risk genes downloaded from the SFARI gene database (https://gene.sfari.org/database/human-gene/) and with a list of 84 high confidence epilepsy risk genes from Wang et al., 2017 (84 epilepsy genes)^63^.

### Immunoprecipitation and western blotting

For immunoprecipitation, cells or brain tissue were lysed in mild lysis buffer (50mM Tris-HCl, 150mM NaCl and 1% NP-40) with cOmplete mini protease inhibitor cocktail (Roche) and centrifuged at 12 000g for 10 minutes at 4 °C. Protein G Dynabeads (Invitrogen) were incubated with primary antibody for 2 hours at room temperature on a rotator, followed by incubation with lysate overnight at 4 degrees on a rotator. Immunoprecipitants were eluted in 2X lamelli sample buffer with ß-mercaptoethanol for 10 minutes at 95 °C.

For western blotting, cells or tissue were lysed in mild lysis buffer or RIPA buffer (50mM Tris-HCl, 150mM NaCl, 0.5% sodium deoxycholate, 0.1% SDS, and 1% NP-40). Samples were loaded into Tris-Glycine gels and transferred to a PVDF membrane (Bio-Rad). Membranes were blocked for 1 hr in 5% milk in 1X TBST, incubated with primary antibody overnight at 4 degrees C, then with secondary antibody (donkey anti-mouse or antirabbit HRP, GE Healthcare) for 1 hr at room temperature before exposure using a ChemiDoc MP system (Bio-Rad).

### TUBE Assay

PNI day 7 iNeurons (without glia) were lysed in TUBE lysis buffer [5nM Tris HCl, 0.15M NaCal, 1mM EDTA, 1% NP40, 10% glycerol, cOmplete mini protease inhibitor cocktail (Roche), 50 µM pr-619 (LifeSensors), 1X 1,10-phenanthroline (LifeSensors)]. 850 µg protein was incubated with 80 µl equilibrated TUBE1 magnetic beads (LifeSensors) at 4 degrees Celsius for 2 hours. Beads were then washed 3X in TBST, resuspended in 1x lamelli sample buffer with ß-mercaptoethanol and boiled for 8 minutes at 95 degrees Celsius. Samples were centrifuged and transferred to a new tube and used for western blotting.

### Cycloheximide-Chase Assay

PNI Day 7 iNeurons (without glia) were treated with cycloheximide solution (20 µg/mL, Sigma). Cells were lysed at 0hrs (untreated), 2 hrs, 5hrs and 9hrs post-treatment in mild lysis buffer and used for western blotting.

### Immunocytochemistry

Cells on glass coverslips were fixed with 4% paraformaldehyde in PBS for 20 min at room temperature. Cells were washed in PBS three times, followed by blocking in blocking/permeabilization solution consisting of 10% Donkey Serum (Millipore) or 1% BSA and 0.3% Triton X-100 (Fisher Scientific) in PBS for 1 hr at room temperature. Incubation in primary antibodies was performed at 4 °C overnight. Cells were then washed in PBS three times, followed by incubation with secondary antibodies in 50% blocking/permeabilization solution at room temperature for 1.5 hr. Cells were then washed in PBS and were mounted on VistaVision glass microscope slides (VWR) using Prolong Gold antifade reagent (Life Technologies).

### Confocal microscopy and morphological analyses

For mouse neuron analyses, confocal images were taken on a Zeiss LSM 700 at a resolution of 2048 x 2048 pixels in an area of 145.16 x 145.16 µm and were processed and analyzed with ImageJ 1.44 software. Sholl analysis was performed using the Sholl analysis plugin in ImageJ. This plugin was used to make concentric circles increasing at a constant radius of 10 µm and to count the number of dendritic intersections. Spine density was calculated by visually counting all protrusions from a dendrite within a 15–25 µm distance starting at a secondary branch point. One to three dendritic segments were analyzed per neuron and the average spine density per neuron was used for statistical analysis. Maximal spine head width (HW), neck width (NW), length (L), and neck length (NL) were measured for each dendritic protrusion using the segmented line tool in ImageJ. Spines were defined as follows: stubby (L < 1 mm), mushroom (1≤ L ≤ 5 mm; HW ≥ 2 x NW), thin (1≤ L ≤ 5 mm; HW ≤ 2 x NW), or filopodia (1≤ L ≤ 5 mm; No visible head). The density of each spine type was calculated by dividing the number of spines in each spine category (mushroom, thin, filopodia, and stubby) by the area of the region of interest (ROI). For axonal morphology analyses on transfected human iNeurons, morphometric analyses were performed of randomly sampled transfected neurons. Images of neurons were acquired using the Zeiss-880 confocal with AIRYSCAN at 20x magnification with 0.6 zoom 1 Airy unit pinhole, Z stack increments of 3.71µm, and laser gain between 500-700 for all channels. These imaging parameters were kept consistent between transductions. Images were processed using NIH image software (ImageJ) as a maximum projection of a single or tiled image. Tiled images were generated using the stitching tool on ImageJ, and image brightness was increased equally among all images. Individual VENUS positive neurons were identified, and SMI-132 positive axons were traced from the base of the soma along the entire projection using the NeuronJ extension. A branch point was defined as a process that extended orthogonal along the axon that exceeded 20µm in length^64^.

To analyze ankyrin-G intensity in the AIS of immunostained iNeurons, confocal images were obtained with a Nikon C2+ confocal microscope. We took confocal images using the 60x oil-immersion objective (NA = 1.40) as z-series of 8 images (2 x 2 stitched; 392.57 µm x 392.57 µm), taken at 0.4 μm intervals, with 1895 × 1895 pixel resolution. Detector gain and offset were adjusted in the channel of cell fill (Venus) to enhance edge detection. Intensity plot profiles for ankyrin-G in the axon were measured by ImageJ with a single plane which was shown the strongest signal in the AIS. Every 5 µm of ankyrin-G intensity was measured and averaged across neurons to produce average intensity plot profiles ± SEM. Confocal images were taken using the 20x objective (NA = 0.75) as z-series of 8–11 images (2 x 2 stitched; 1185.43 µm x 1185.43 µm), taken at 0.4 μm intervals, with 1895 × 1895 pixel resolution. MAP2 stained images were obtained, and traces of dendrites were drawn and analyzed with Sholl analysis in ImageJ.

### SIM imaging and analysis

Imaging and reconstruction parameters were empirically determined with the assistance of the expertise in the Nikon Imaging Center at Northwestern. The acquisition was set to 10MHz, 14 bit with EM gain, and no binning. Auto exposure was kept between 100-300ms, and the EM gain multiplier restrained below 300. Conversion gain was held at 1x. Laser power was adjusted to keep LUTs within the first quarter of the scale (<4000). Reconstruction parameters (0.96, 1.19, and 0.17) were kept consistent across experiments and imaging sessions. The resolution of images was taken with 2048 x 2048 pixels in the area of 66.56 x 66.56 µm. For each spine analyzed, the single-plane in which the spine head was in focus, based on the cell fill, was chosen for analysis. Three shape classes, mushroom, thin, and stubby, were assigned for analyzing spine morphologies. Spines were determined as mushroom if the diameter of the head was much greater than the diameter of the neck. Spines were determined as thin if serial viewing revealed the length to be greater than the neck diameter, and the diameters of the head and neck to be similar. Spines were determined as stubby if the diameter of the neck was similar to the total length of the spine. Using ImageJ software, each spine head was outlined manually in the channel of the cell fill to detect the area. A specific dendritic region (WT: 59.03±4.96 µm; Df(h15q13)/+: 67.40±4.33 µm; Control (Fam1): 120.97±12.49 µm; 15q13.3 HET (Fam 1): 105.18±7.49 µm; OTUD7A^L233F/L233F^: 100.87±7.28 µM) was selected, and puncta counts were made for the measurement of Ankyrin-G puncta; puncta smaller than 0.065 μm^2^ were excluded from the analysis. Visual assessment of fluorescence intensity was used to delineate separate or connected puncta. Puncta were considered separate if a region of decreased intensity was readily visible. The number of puncta within the spine head was quantified manually and recorded. For the dendritic analysis, the first branched apical dendrite from a pyramidal neuron was outlined automatically by thresholding 10,000 in a 16-bit image. In the chosen region of interest (ROI) of the dendritic shaft without spines, Ankyrin-G puncta were analyzed with the option of analyze particles automatically by ImageJ; also, puncta smaller than 0.065 μm^2^ were excluded. All images were processed with converting mask and watershed with a binary option.

### Statistical Analysis

Data are expressed as mean ± SEM. Blinding was performed for SIM imaging experiments. Outliers were identified with the ROUT method (Q=1%). Normality tests were performed using the D’Agostino & Pearson test, Shapiro-Wilk test and Kolmogorov-Smirnov test. For normally distributed datasets, we used the Student’s t test, one-sample t test, one-way ANOVA with Dunnett’s post-hoc test, two-way ANOVA with Dunnett’s post-hoc test, and Repeated Measures two-way ANOVA with Tukey’s post-hoc test in GraphPad Prism 9 statistical software for statistical analyses. For non-normally distributed data, we used non-parametric statistical tests (Mann-Whitney test or Kruskal-Wallis test). p values in the figure legends are from the specified tests, and p < 0.05 was considered statistically significant. All error bars represent standard error of the mean (SEM). For SAINT analysis, a cutoff SAINT score of 0.6 was used for significance. For functional enrichment analysis, an FDR <0.05 was considered significant.

## ACKNOWLEDGEMENTS

This work was supported by grants to K.K.S., (Canadian Institutes of Health Research [CIHR], Natural Sciences and Engineering Research Council [NSERC], the Ontario Brain Institute (OBI), and The Network of European Funding for Neuroscience Research (ERA-NET Neuron) and P.P. (R01MH097216). We would also like to thank Dr. Vann Bennett (Duke University) for his advice related to Ankyrin-G experiments and reagents. All schematics were created with BioRender.com.

## AUTHOR CONTRIBUTIONS

This study was designed by KKS and BKU. The majority of the experiments were performed by BKU, with assistance from SY, LC, SK, SX, YL, NM, AA and AC. BKU developed and performed the BioID2 experiments, MEAs and morphological analysis of primary mouse cortical neurons, lentivirus production, biochemical experiments, and all genetic rescue experiments. LC grew and induced iPSC-derived iNeurons and performed MEA and patch-clamp electrophysiology on iNeurons. SK contributed to co-immunoprecipitation and axon morphology experiments. SY performed SIM imaging and AIS Ankyrin-G analysis. AA reprogrammed Family 3 iPSCs. Liquid Chromatography and Tandem Mass Spectrometry (LC-MS/MS) for BioID2 was performed by SX and YL. AC performed flow cytometry for lentiviral titering. EM, GR, JH, LF, AV, ACE, AV and SWS performed patient recruitment, consent and sample collection. BWD and NM contributed to study design. PP provided HA-Ankyrin-G domain plasmids and supervised SY. Data were analyzed and figures prepared by BU. BU and KKS wrote the manuscript.

## DATA AVAILABILITY

Raw proteomics data from BioID2 experiments will be uploaded to the ProteomeXchange (http://www.proteomexchange.org).

## SUPPLEMENTARY MATERIAL

Supplementary Figures 1-10 are attached as a separate pdf file. Supplementary Tables are attached as separate excel files.

**Supplementary Table 1.** Human iNeuron cellular phenotyping

**Supplementary Table 2.** SAINTexpress analysis tables

**Supplementary Table 3.** Significant BioID2 hits

**Supplementary Table 4.** gProfiler tables

**Supplementary Table 5.** OTUD7A BioID2 ASD-associated SFARI genes and Epilepsy genes

**Supplementary Table 6.** OTUD7A BioID2 mutation analysis

