## Supplementary Figures for "Impaired OTUD7A-dependent Ankyrin regulation mediates neuronal dysfunction in mouse and human models of the 15q13.3 microdeletion syndrome"

### Supplementary Figure 1

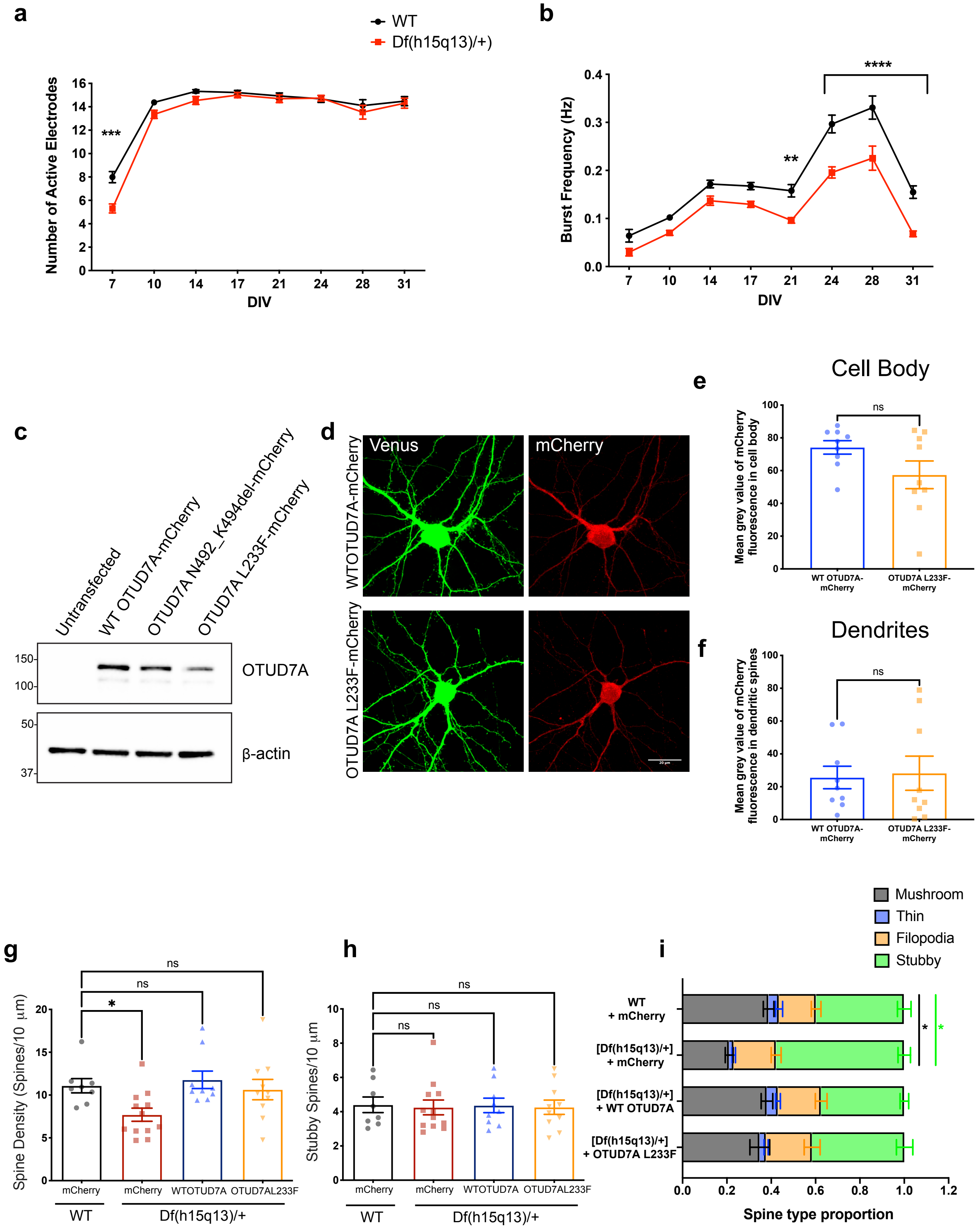

**Supplementary Figure 1. Additional MEA analyses and genetic rescue of morphological deficits in *Df(h15q13)/+* cortical neurons**

**(a)** The average number of active electrodes per well from MEA recordings of WT and *Df(h15q13)/+* cortical neurons. n=77 wells WT, 65 wells Df(h15q13)/+ from 3 mouse cortical cultures on 3 MEA plates. Multiple Mann Whitney tests with two-stage step-up method (Benjamini, Krieger, and Yekutieli), ***$q=0.000894$, $U=1582$, Mean rank of WT= 83.45, Mean rank of Df(h15q13)/+=57.34.
**(b)** Burst frequency analysis. n=77 wells WT, 65 wells Df(h15q13)/+ from 3 mouse cortical cultures on 3 MEA plates. Repeated Measures Two-Way ANOVA with Sidak's post-hoc test. *$p<0.05$, **$p<0.01$, ***$p<0.001$, ***$p<0.0001$. Interaction: $F(7,980)=3.608$, $p=0.0008$; DIV: $F(3.408,477.1)=87.57$, $p<0.0.0001$; Genotype: $F(1,140)=36.59$, $p<0.0001$; Subject: $F(140,980)=3.00$,$p<0.0001$.
**(c)** Validation of WT and mutant OTUD7A-mCherry construct protein expression in transfected HEK293 FT cells. Anti-OTUD7A antibody detected a band at the expected size in all transfected samples. Only WT OTUD7A-mCherry and OTUD7A L233F-mCherry were used for further re-expression experiments.
**(d)** Representative confocal images of co-transfected WT DIV 14 mouse cortical neurons. Cells were stained with mCherry antibody. 63X objective, scale bar=20 μm.
**(e-f)** Comparison of WT and mutant OTUD7A-mCherry tagged expression levels in primary mouse cortical neurons. No significant differences in expression were observed in the cell body **(e)** (Student's t-test:$p=0.0939$, $t=1.781$, $df=16$) or dendrites **(f)** (Student's t-test: $p=0.8360$, $t=0.2104$, $df=16$) n= 9 neurons per condition.
**(g)** Spine density analysis of co-transfected WT and *Df(h15q13)/+* cortical neurons. n= 8 neurons WT + mCherry, 12 neurons [Df(h15q13)/+] + mCherry, 9 neurons [Df(h15q13)/+] + WT OTUD7A-mCherry, 10 neurons [Df(h15q13)/+] + OTUD7A L233F. Samples were taken from 3 mouse cultures. *$p<0.05$, Kruskal Wallis test with Dunn's post-hoc test. Kruskal-Wallis statistic=11.51, $p=0.0093$ (Approximate).
**(h)** Stubby spine density. n= 8 neurons WT + mCherry, 12 neurons [Df(h15q13)/+] + mCherry, 9 neurons [Df(h15q13)/+] + WT OTUD7A-mCherry, 10 neurons [Df(h15q13)/+] + OTUD7A L233F. Samples were taken from 3 mouse cultures. One-Way ANOVA with Dunnett's post-hoc test. $F(3,35) = 0.02962$, $p=0.9330$.
**(i)** Analysis of spine type proportions. n= 8 neurons WT + mCherry, 12 neurons [Df(h15q13)/+] + mCherry, 9 neurons [Df(h15q13)/+] + WT OTUD7A-mCherry, 10 neurons [Df(h15q13)/+] + OTUD7A L233F. Samples were taken from 3 mouse cultures. Two-Way ANOVA with Dunnett's post-hoc test. ****$p<0.0001$, Interaction: $F(9,140) = 8.810$, $p<0.0001$; Spine Type: $F(3,140) = 181.8$, $p<0.0001$; Condition: $F(3,140) = 0.0003015$, $p>0.9999$.

#### Supplementary Figure 2

**a**

| Family | Status | Sex | CNV or mutation | Genes Involved | Known NDDs | Family History |
|---|---|---|---|---|---|---|
| Family 1 | Mother (Control) | F | — | — | — | Unknown |
| | Proband | F | 15q13.2-13.3 (BP4-BP5) | ARHGAP11B, FAN1, MTMR10, TRPM1, MIR211,KLF13, OTUD7A, CHRNA7 | ASD | |
| OTUD7A L233F | Mother (Carrier) | M | chr15:g.31819467G>A NM_130901.2: c.697C>T | OTUD7A | — | Consanguineous parents, brother had nonspecific difficulties, both parents had learnng disability |
| | Father (Carrier) | F | chr15:g.31819467G>A NM_130901.2: c.697C>T | OTUD7A | — | |
| | Proband | M | chr15:g.31819467G>A NM_130901.2: c.697C>T (homozygous) | OTUD7A | Severe global developmental delay, language impairment and epileptic encephalopathy | |
| | Brother (Carrier) | M | chr15:g.31819467G>A NM_130901.2: c.697C>T | OTUD7A | — | |
| Family 2 | Mother (Carrier) | F | 15q13.2-13.3 (BP4-BP5) | FAN1, MTMR10, TRPM1, MIR211, KLF13, OTUD7A,CHRNA7 | — | Two paternal aunts with seizure disorders, both parents had comprehension/ learning difficulties |
| | Proband | F | 15q13.2-13.3 (BP4-BP5) | FAN1, MTMR10, TRPM1, MIR211, KLF13, OTUD7A, CHRNA7 | Absence seizures, DD, ID, Learning disorder | |
| | Brother (Control) | M | — | — | — | |
| Family 3 | Father (Control) | M | — | — | — | Unknown |
| | Proband | F | 15q13.2-13.3 (BP4-BP5) | ARHGAP11B, FAN1, MTMR10, TRPM1, MIR211, KLF13, OTUD7A,CHRNA7 | Asperger's (ASD), ADHD, absence epilepsy | |

**b**

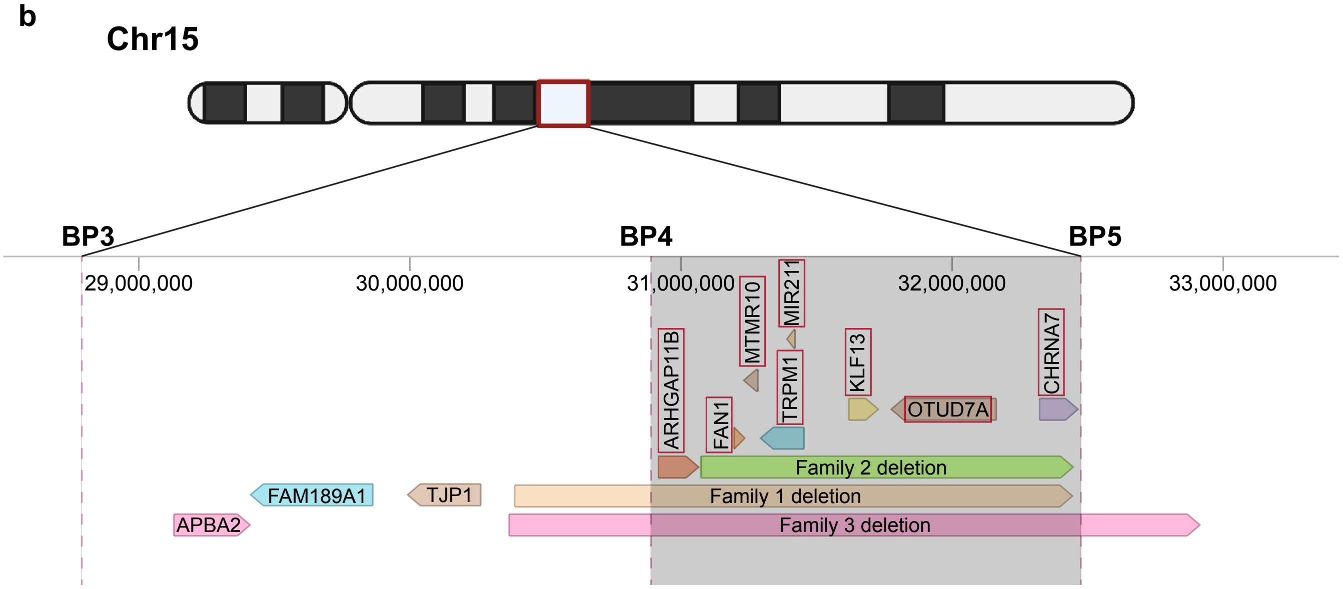

**Supplementary Figure 2. Genetic and clinical information for human iPSC lines**

**(a)** Clinical information associated with patient and familial control lines. The OTUD7A L233F Family carriers (maternal, paternal and brother) were not used in this study.
**(b)** Schematic of chromosome 15 showing the 15q13.3 BP3,BP4 and BP5 deletion breakpoints and genes involved.

#### Supplementary Figure 3

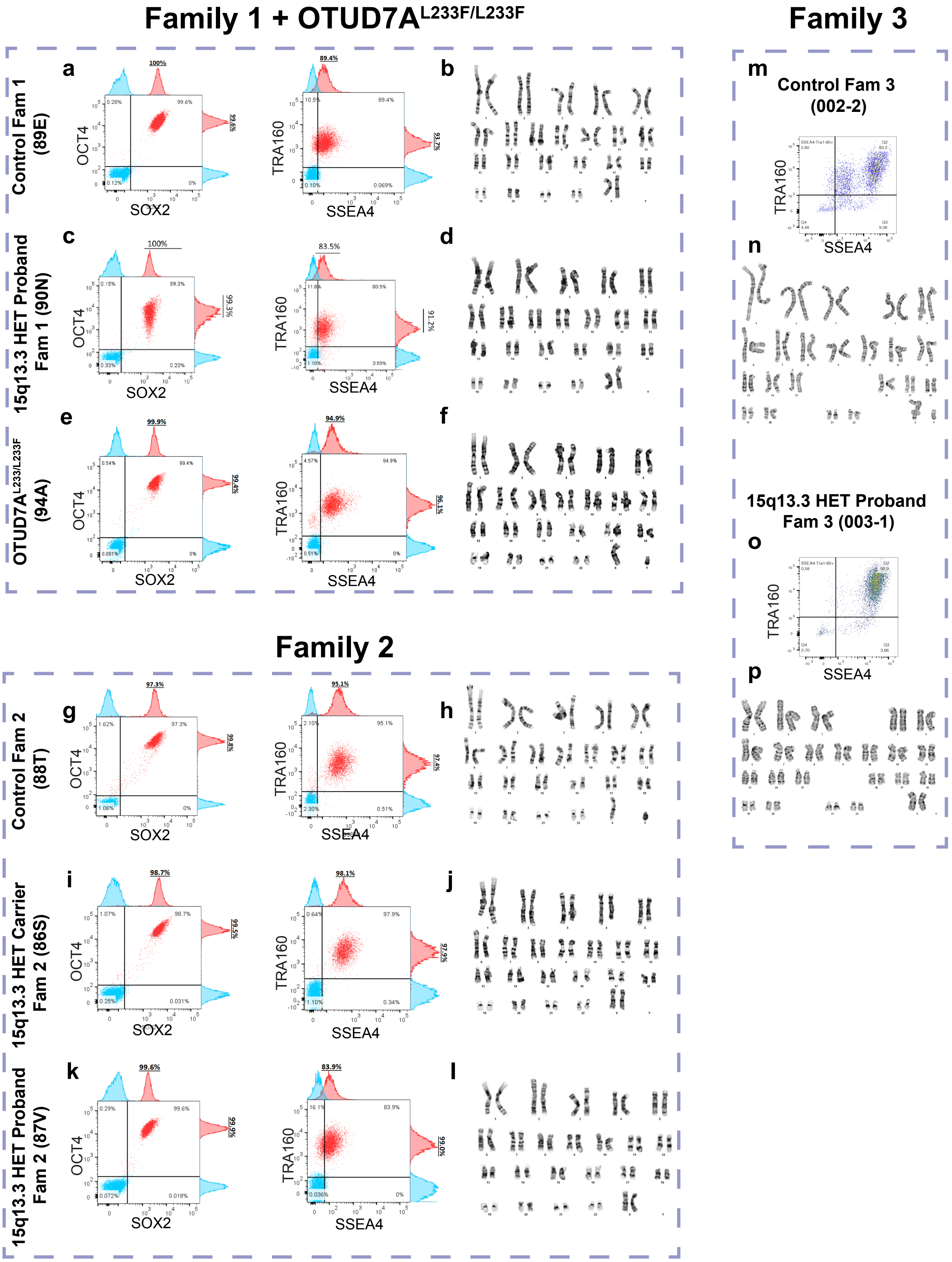

**Supplementary Figure 3. iPSC reprogramming validation of human patient samples**

Flow cytometry plots of cell populations expressing pluripotency markers in hiPSCs from **(a)** Control (Fam 1), **(c)** 15q13.3 HET (Fam 1) and the **(e)** OTUD7AL233F/L233F patient. Left: OCT4/SOX2 Right: TRA160/SSEA4.
G-banding analysis shows normal karyotype in hiPSCs from **(b)** Control (Fam 1), **(d)** 15q13.3 HET (Fam 1) and the **(f)** OTUD7AL233F/L233F patient.
Flow cytometry plots of cell populations expressing pluripotency markers in hiPSCs from **(g)** Control (Fam 2), **(i)** 15q13.3 HET Carrier (Fam 2) and **(k)** 15q13.3 HET proband (Fam 2) Left: OCT4/SOX2 Right: TRA160/SSEA4.
G-banding analysis shows normal karyotype in hiPSCs from **(h)** Control (Fam 2), **(j)** 15q13.3 HET Carrier (Fam 2) and **(l)** 15q13.3 HET proband (Fam 2).
Flow cytometry plots of cell populations expressing pluripotency markers (TRA160/SSEA4) in hiPSCs from **(m)** Control (Fam 3) and **(o)** 15q13.3 HET proband (Fam 3).
G-banding analysis shows normal karyotype in hiPSCs from **(n)** Control (Fam 3) and **(p)** 15q13.3 HET proband (Fam 3).

### Supplementary Figure 4

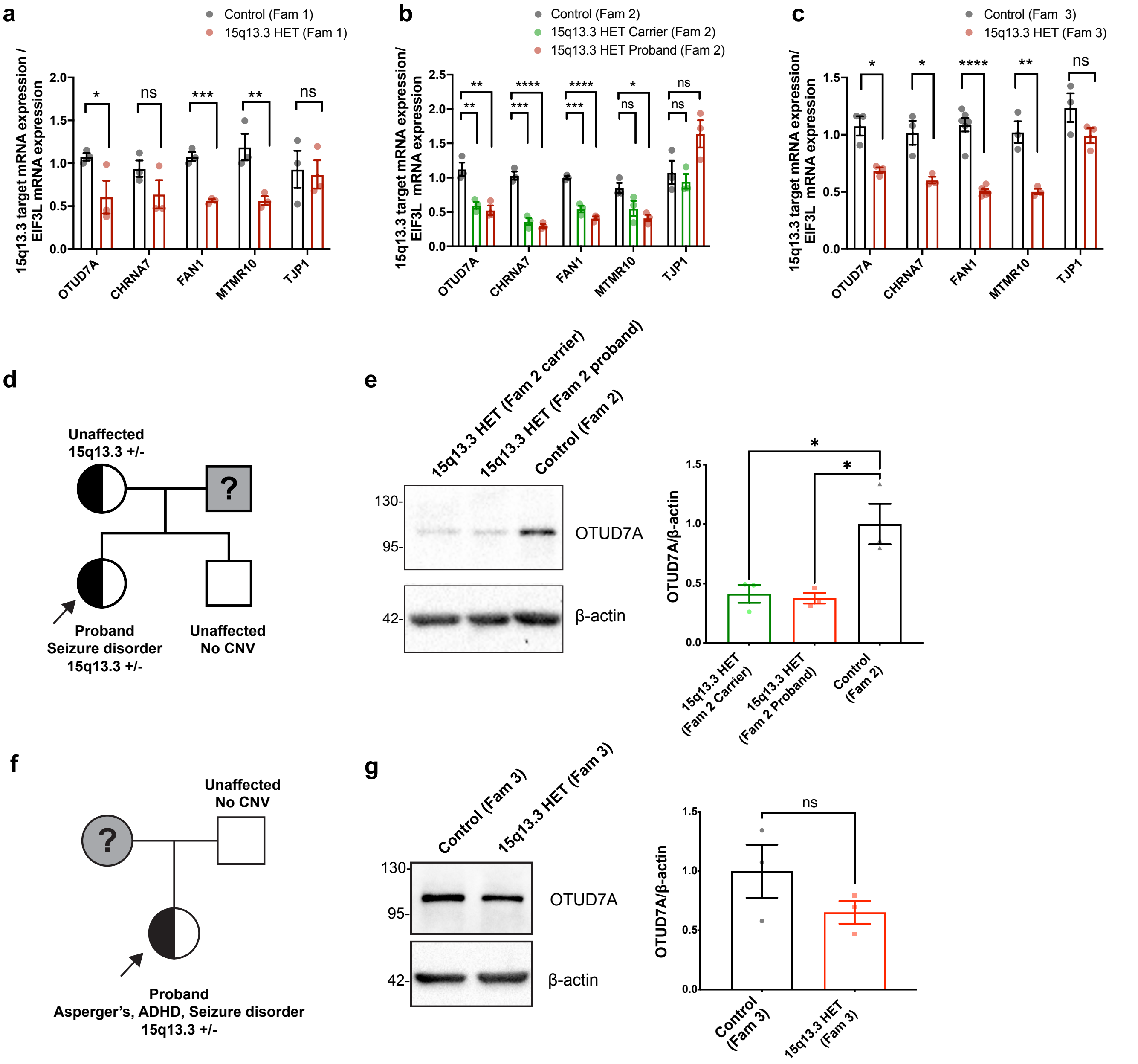

**Supplementary Figure 4. mRNA and protein expression levels of 15q13.3 microdeletion genes in hIPSC-derived iNeurons**

**(a-c)** mRNA expression of 15q13.3 microdeletion genes and breakpoint flanking gene (TJP1) relative to EIF3L in iNeurons derived from **(a)** Family 1, **(b)** Family 2, and **(c)** Family 3. N= 3 Ngn2/Rtta transductions per line. Unpaired two-tailed t-tests, *p<0.05, **p<0.01, ***p<0.001, ****p<0.000
**(d)** Family 2 pedigree.
**(e)** Representative western blot (left) and quantification (right) of OTUD7A levels in Family 2 iNeurons; n= 3 separate Ngn2/Rtta transductions per line; One-Way ANOVA with Dunnett's post-hoc test, *p<0.05, F(2.6)=7.921, p=0.0121.
**(f)** Family 3 pedigree.
**(g)** Representative western blot (left) and quantification (right) of OTUD7A levels in Family 3 iNeurons; n= 3 separate Ngn2/Rtta transductions per line; Unpaired two-tailed t-test, p=0.2279, t=1.423, df=4.

### Supplementary Figure 5

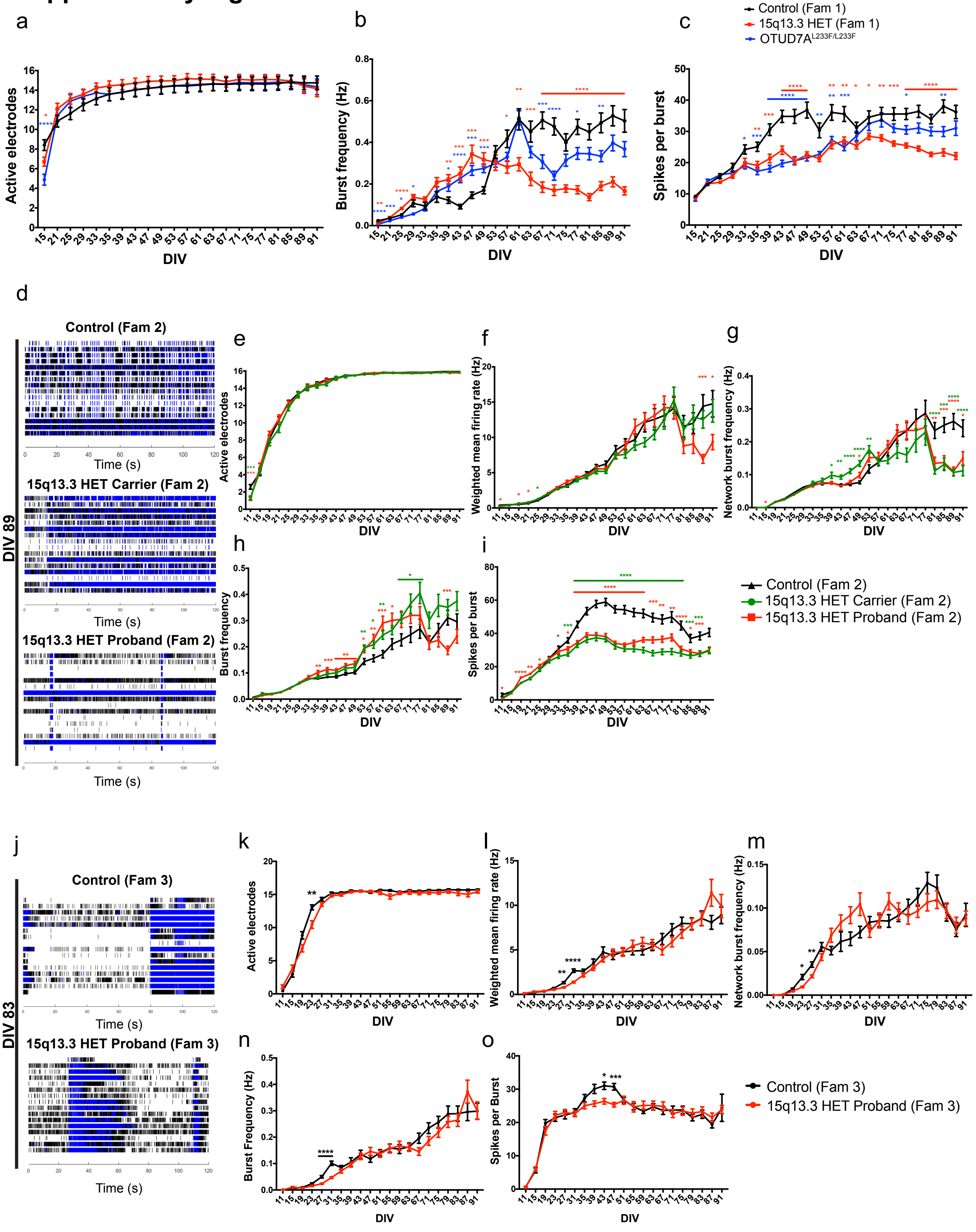

**Supplementary Figure 5. Additional MEA analysis parameters in hIPSC-derived iNeurons.**

**(a)** Number of active electrodes (Interaction: F(42, 1953)=4.137, p<0.0001; p<0.0001; DIV: F(3.906,363.3) = 182.2, p<0.0001; Genotype: F(F(2,93)=0.2100, p=0.8110; Subject: F(93,1953) = 142.2, p<0.0001)
**(b)** Burst frequency (Interaction: F(42,1848) = 17.64, p<0.0001; DIV: F(5.532,486.8) = 84.47, p<0.0001; Genotype: F(2,88) =9.998, p=0.0001; Subject: F(88,1848) = 11.38, p<0.0001)
**(c)** Spikes per burst (Interaction: F(42,1848)=8.060, p<0.0001; DIV: F(6.108, 537.5) = 110.9, p<0.0001; Genotype: F(2,88) = 19.08, p<0.0001; Subject: F(88,1848) = 1824, p<0.0001) analyzed from MEA recordings of Family 1 and OTUD7A$^{L233F/L233F}$ patient iNeurons. Control (Fam 1) n=29 wells, 15q13.3 HET (Fam 1) n=29 wells, OTUD7A$^{L233F/L233F}$ n=30 wells from two separate NGN2/Rtta transductions. Repeated Measures Two-Way ANOVA with Dunnett's post-hoc test, *p<0.05, **p<0.01, ***p<0.001, ***p<0.0001.
**(d)** Raster plots of MEA recordings of neural network activity from DIV 89 Family 2 human iNeurons.
**(e)** The number of active electrodes (Interaction: F (44, 2750) = 1.803, P=0.0010; DIV: F (4.733, 591.7) = 1438, P<0.0001, Genotype: F (2, 125) = 0.742, P=0.4781, Subject: F (125, 2750) = 10.82, P<0.0001),
**(f)** Weighted mean firing rate (Interaction: F (44, 2750) = 2.730, P<0.0001, DIV: F (4.468, 558.5) = 115.4, P<0.0001, Genotype: F (2, 125) = 0.6183,P=0.5405, Subject: F (125, 2750) = 8.261, P<0.0001),
**(g)** Network burst frequency (Interaction: F (44, 2750) = 5.186, P<0.0001; DIV: F (5.475, 684.4) = 79.13, P<0.0001; Genotype: F (2, 125) = 3.815, P=0.0246; Subject: F (125, 2750) = 4.982, P<0.0001),
**(h)** Burst frequency (Interaction: F (44, 2750) = 4.436, P<0.0001; DIV: F (5.747, 718.4) = 123.6, P<0.0001; Genotype: F (2, 125) = 6.256, P=0.0026; Subject: F (125, 2750) = 5.100", P<0.0001)
**(i)** Spikes per burst Interaction: F (44, 2772) = 18.09, P<0.0001; DIV: F (5.025, 633.1) = 348.5, P<0.0001; Genotype: F (2, 126) = 31.98, P<0.0001; Subject: F (126, 2772) = 22.80, P<0.0001) analyzed from MEA recordings of Family 2 iNeurons. Control (Fam 2) n= 42 wells, 15q13.3 HET Carrier (Fam 2) n=42 wells, 15q13.3 HET Proband (Fam 2) n=44 wells from 3 separate Ngn2/Rtta transductions. Repeated Measures Two-Way ANOVA with Dunnett's post-hoc test, *p<0.05, **p<0.01, ***p<0.001, ***p<0.0001.
**(j)** Raster plots of MEA recordings of neural network activity from DIV 87 Family 3 human iNeurons.
**(k)** The number of active electrodes (Interaction: F (20, 1620) = 3.941, P<0.0001; DIV: F (4.325, 350.3) = 505.2, P<0.0001; Genotype: F (1, 81) = 5.408, P=0.0225; Subject: F (81, 1620) = 4.989, P<0.0001),
**(l)** Weighted mean firing rate (Interaction: F (20, 1620) = 1.970, P=0.0064; DIV: F (3.143, 254.6) = 72.16, P<0.0001; Genotype: F (1, 81) = 0.009191, P=0.9239; Subject: F (81, 1620) = 8.044, P<0.0001),
**(m)** Network burst frequency (Interaction: F (20, 1620) = 1.989, P=0.0057; DIV: F (7.752, 627.9) = 48.33, P<0.0001; Genotype: F (1, 81) = 0.03251, P=0.8574; Subject: F (81, 1620) = 3.764, P<0.0001),
**(n)** Burst frequency (Interaction: F (20, 1620) = 1.779, P=0.0181; DIV: F (3.531, 286.0) = 86.34,P<0.0001; Genotype: F (1, 81) = 0.4747, P=0.4928; Subject: F (81, 1620) = 8.403, P<0.0001), and
**(o)** Spikes per burst (Interaction: F (20, 1620) = 1.763, P=0.0197; DIV: F (4.995, 404.6) = 73.61, P<0.0001; Genotype: F (1, 81) = 0.4789, P=0.4909; Subject: F (81, 1620) = 5.614, P<0.0001) analyzed from MEA recordings of Family 3 iNeurons. Control (Fam 3) n= 35 wells, 15q13.3 HET Proband (Fam 3) n=48 wells from 3 separate Ngn2/Rtta transductions. Repeated Measures Two-Way ANOVA with Sidak's post-hoc test, *p<0.05, **p<0.01, ***p<0.001, ***p<0.0001.

### Supplementary Figure 6

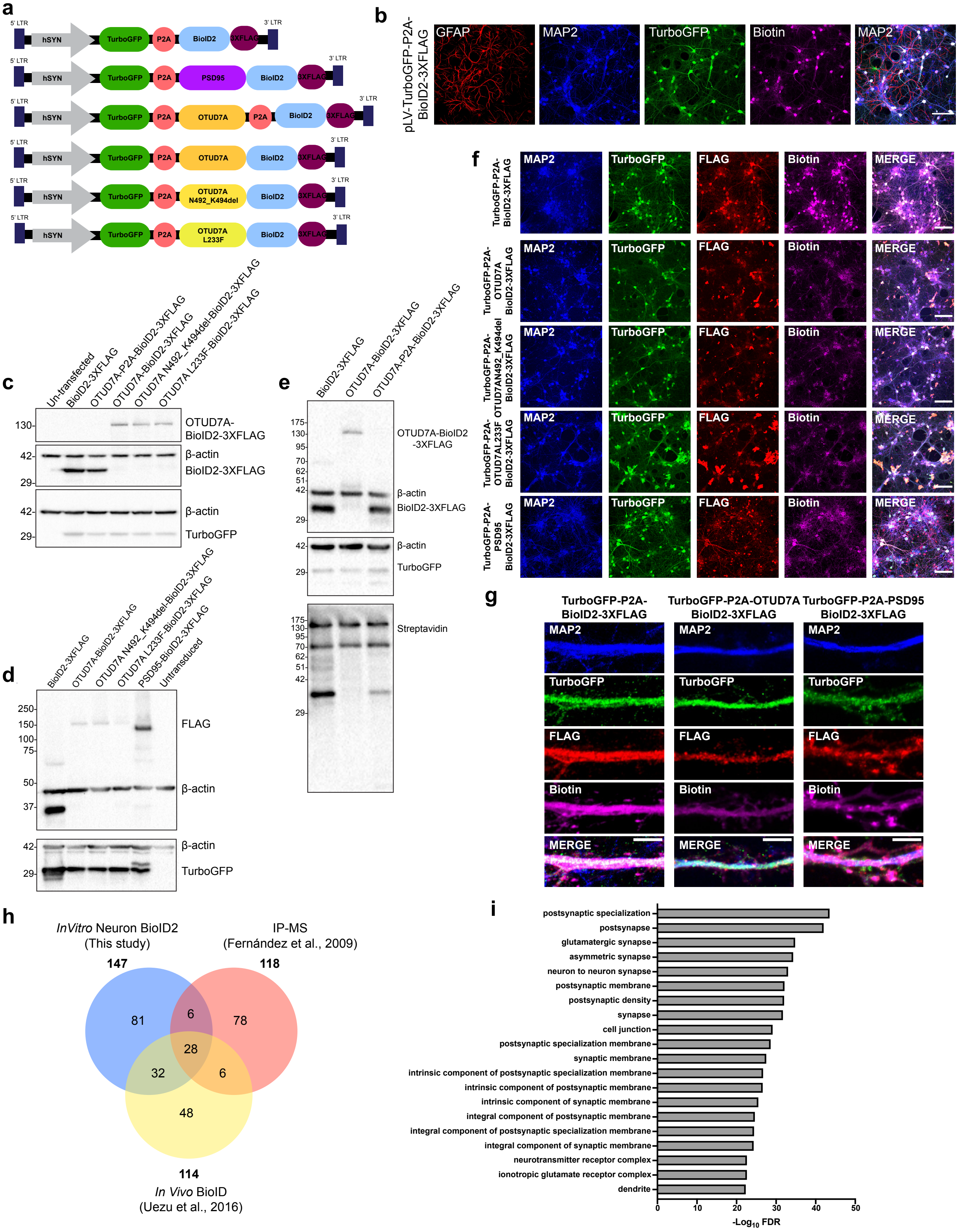

**Supplementary Figure 6. Validation of neuron-specific BioID2 system**

**(a)** Schematic of the lentiviral BioID2 fusion constructs used in the study.
**(b)** Confocal image of DIV 18 mouse cortical neuron + glia co-culture transduced with hSYN-TurboGFP-P2A-BioID2-3XFLAG lentivirus and stained for GFAP, MAP2, TurboGFP and Biotin. The BioID2 construct shows neuron-specific expression. Objective 20X, Scale bar= 100 µm.
**(c)** Western blot of protein lysates from DIV 18 CD1 mouse cortical neurons transduced with the indicated BioID2 constructs at DIV 14 (MOI=0.9).
**(d)** Western blot of protein lysates from HEK293FT cells transfected with the indicated BioID2 constructs.
**(e)** Western blot using protein lysates from primary cortical neurons were transduced at DIV 5, followed by 50 µM biotin treatment at DIV 8. Cells were then lysed and probed with antibodies against, Flag, ß-actin, TurboGFP and HRP-conjugated streptavidin antibody.
**(f)** and **(g)** Primary cortical neurons were transduced at DIV 14, followed by 50 µM biotin treatment at DIV 17. Cells were fixed at DIV 18 and stained with antibodies against TurboGFP, FLAG, Streptavidin and MAP2. **(i)** Objective 20X, Scale Bar: 100µm; **(j)** Objective 63X, Scale Bar:5µm.
**(h)** Comparison of PSD95-BioID2 interactors in this study with PSD95 interactors identified in Uezu et al., 2016 (*In Vivo* AAV PSD95-BioID) and in Fernandez et al., 2009 (IP-MS).
**(i)** Top 20 GO: Cellular component pathways enriched in the PSD95-BioID2 dataset. Functional enrichment analysis was performed using gProfiler with Bonferroni correction for multiple testing. A custom background statistical domain scope was used (Sharma et al., 2015, mouse whole brain proteome).

### Supplementary Figure 7

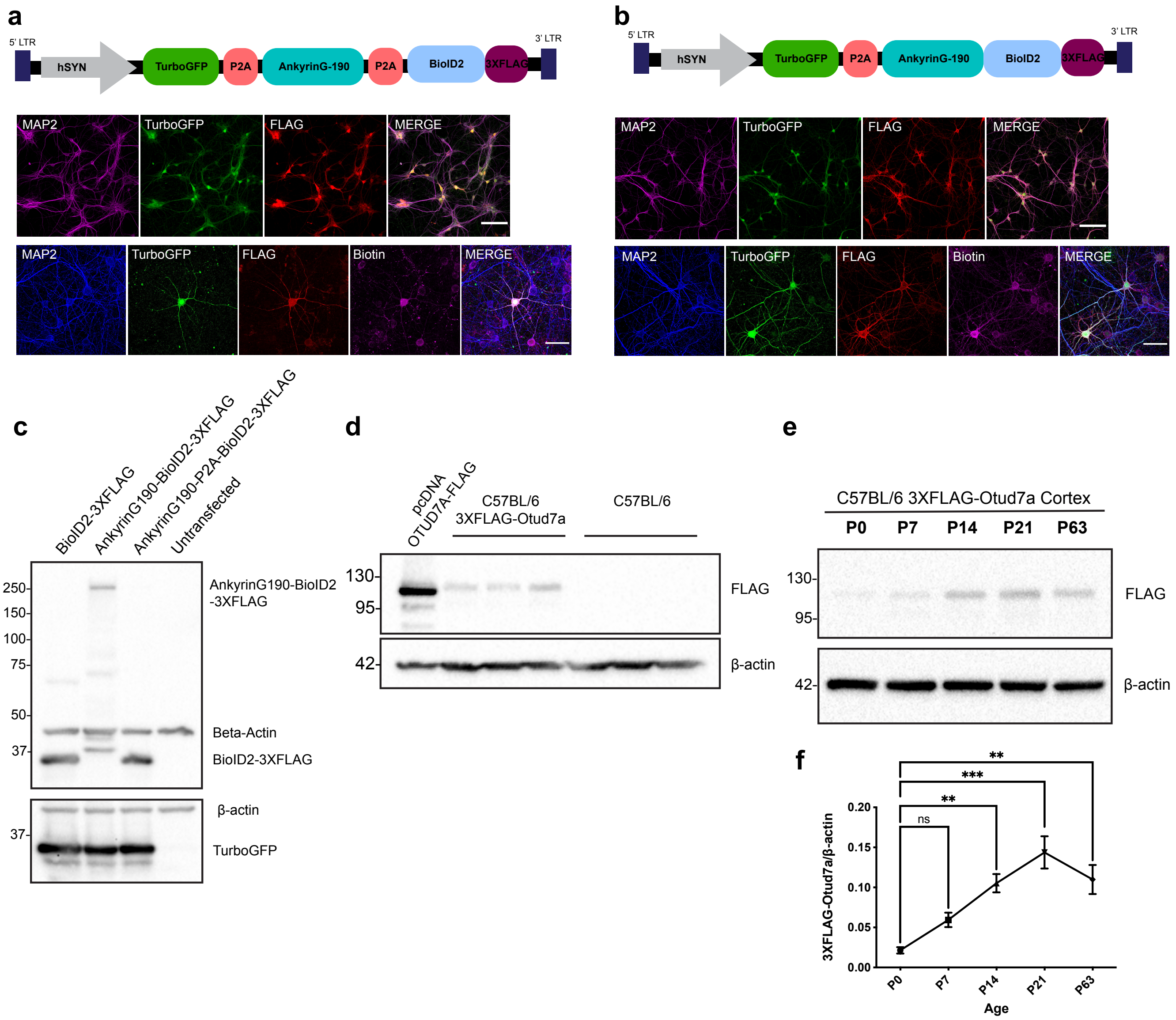

**Supplementary Figure 7. Validation of Ankyrin-G-190 BioID2 constructs and C57BL/6-3XFLAG-Otud7a mice**

**(a)** Top: Schematic of Ankyrin-G-190-P2A-BioID2-3XFLAG control lentiviral construct.
Middle: Confocal image of CD1 WT mouse cortical neurons transduced with Ankyrin-G-190-P2A-BioID2-3XFLAG at DIV 5 and fixed at DIV 9. Cells were stained with antibodies against MAP2, TurboGFP, and FLAG Objective 20X, Scale bar = 100
Bottom: Confocal image of CD1 WT mouse cortical neurons transduced with Ankyrin-G-190-P2A-BioID2-3XFLAG at DIV 14 and fixed at DIV 18. Cells were stained with antibodies against MAP2, TurboGFP, FLAG and Biotin. Objective 40X, Scale bar = 50 µm.
**(b)** Top: Schematic of Ankyrin-G-190-BioID2-3XFLAG lentiviral construct.
Middle: Confocal image of CD1 WT mouse cortical neurons transduced with Ankyrin-G-190-BioID2-3XFLAG at DIV 5 and fixed at DIV 9. Cells were stained with antibodies against MAP2, TurboGFP, and FLAG Objective 20X, Scale bar = 100 µm.
Bottom: Confocal image of CD1 WT mouse cortical neurons transduced with Ankyrin-G-190-BioID2-3XFLAG at DIV 14 and fixed at DIV 18. Cells were stained with antibodies against MAP2, TurboGFP, FLAG and Biotin. Objective 40X, Scale bar = 50 µm.
**(c)** Validation of BioID2 constructs used in the Ankyrin-G-190 BioID2 experiment. Western blot on protein lysates from HEK 293 FT cells transfected with the indicated constructs.
**(d)** Western blot on lysates from P20 C57BL/6-3XFLAG-Otud7a and C57BL/6J mouse cortex and HEK293FT cells transfected with pcDNA-OTUD7A-FLAG.
**(e)** Representative western blot from C57BL/6-3XFLAG-Otud7a mouse cortex harvested at the indicated ages.
**(f)** 3XFLAG-Otud7a levels are significantly increased at P14, P21 and P63 compared to P0. N= 3 cortices per condition per time-point, $**p<0.01$, $***p<0.001$, One-Way ANOVA with Dunnett's post-hoc test, $F(4, 10) = 11.85$, $P=0.0008$.

#### Supplementary Figure 8

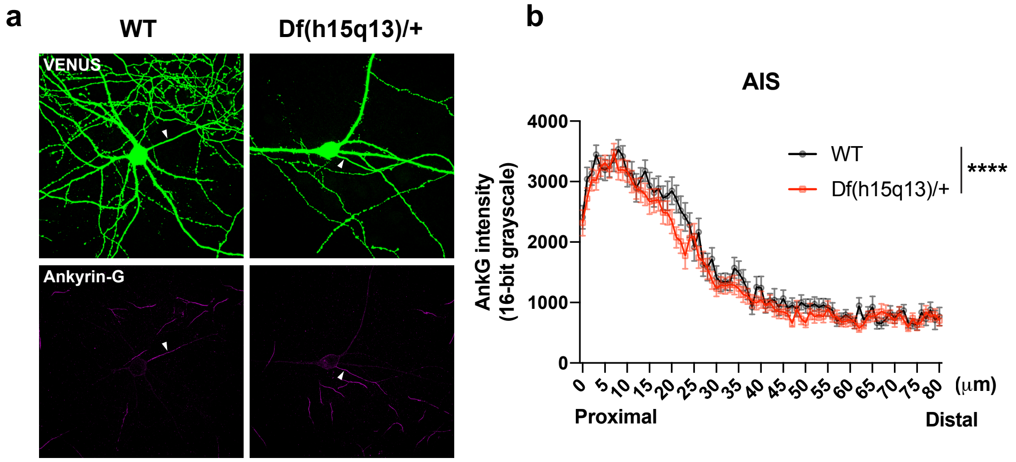


**Supplementary Figure 8. Analysis of AIS Ankyrin-G levels in WT and *Df(h15q13)/+* mouse neurons**

**(a)** Representative confocal images of WT and *Df(h15q13)/+* neurons transfected with VENUS and stained for Ankyrin-G. Scale bar = 50 µm. Arrow indicates location of AIS of transfected neuron.
**(b)** Quantification of Ankyrin-G intensity (mean grey value) at the AIS. WT: n = 32 neurons, *Df(h15q13)/+*: n= 42 neurons. Two-Way ANOVA with Sidak's post-hoc test; Interaction: $F (80, 5772) = 0.6872$, $P=0.9849$; Distance from soma: $F (80, 5772) = 78.36$, $P<0.0001$; Genotype: $F (1, 5772) = 30.69$, $P<0.0001$.

#### Supplementary Figure 9

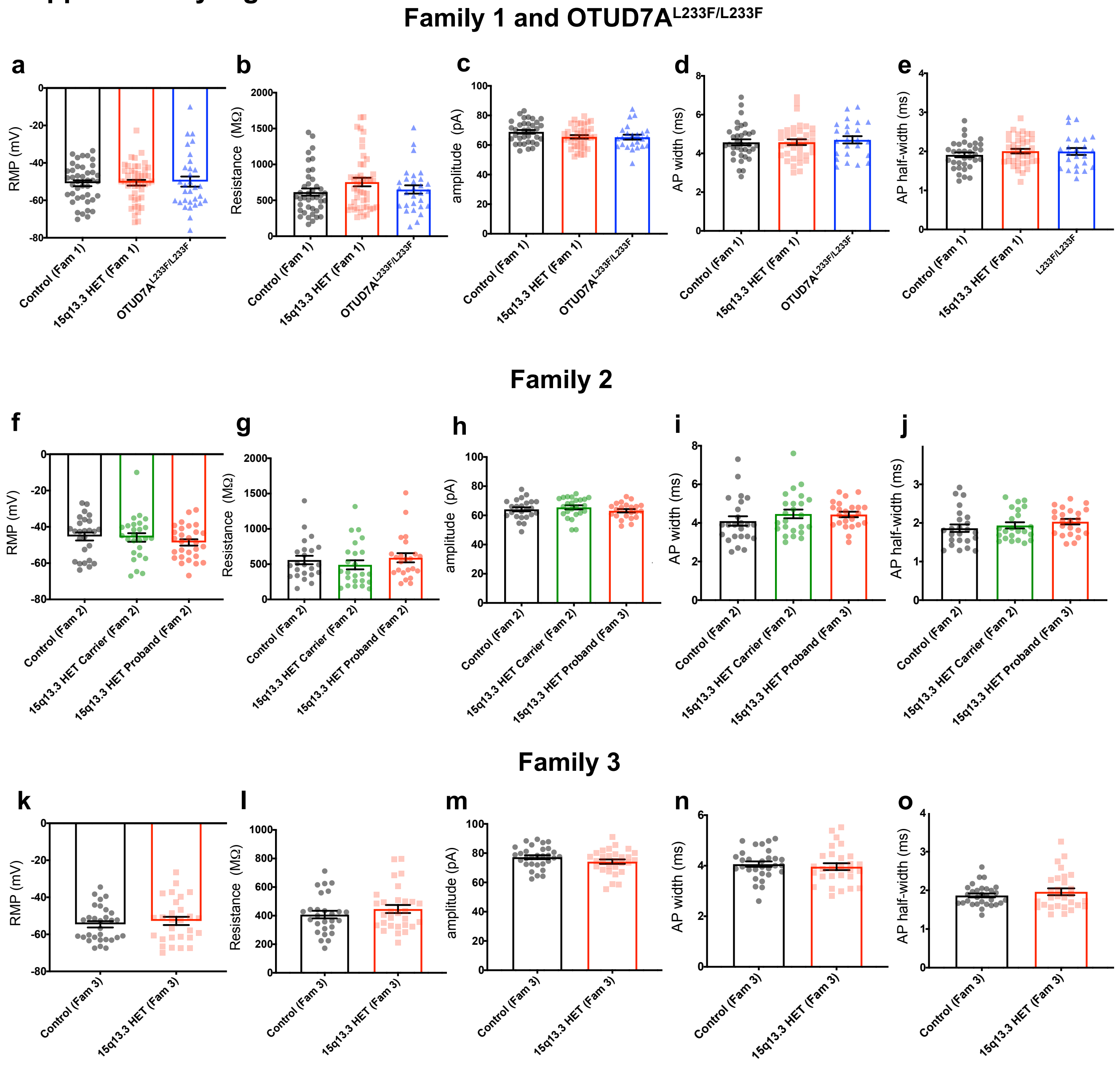

**Supplementary Figure 9. Additional intrinsic membrane and action potential properties in hiPSC-derived iNeurons.**

**(a-e)** Intrinsic electrophysiological properties of Family 1 and OTUD7A$^{L233F/L233F}$ iNeurons.
**(a)** Resting membrane potential (One-Way ANOVA with Dunnett's post-hoc test, F (2, 115) = 0.05298, P=0.9484; Control n= 42, 15q13.3 HET n=45, OTUD7A$^{L233F/L233F}$ n=31).
**(b)** Membrane resistance (Kruskal-Wallis test with Dunn's post-hoc test, Kruskal-Wallis statistic= 3.014, p=0.2215, Control n= 41, 15q13.3 HET n=44, OTUD7A$^{L233F/L233F}$ n=29)
**(c)** Action potential amplitude from threshold (One-Way ANOVA with Dunnett's post-hoc test; F (2, 94) = 2.391, P=0.0971; Control n= 35, 15q 13.3 HET n=38, OTUD7A$^{L233F/L233F}$ n=24.
**(d)** Action potential width (One-Way ANOVA with Dunett's post-hoc test, F (2, 93) = 0.1701, P=0.8439; Control n= 35, 15q13.3 HET n=37, OTUD7A$^{L233F/L233F}$ n=24)
**(e)** Action potential half-width (One-Way ANOVA with Dunnett's post-hoc test, F (2, 93) = 0.6471, P=0.5259; Control n= 34, 15q13.3 HET n=38, OTUD7A$^{L233F/L233F}$ n=24)
**(f-j)** Intrinsic electrophysiological properties of Family 2 iNeurons.
**(f)** Resting membrane potential (One-Way ANOVA with Tukey's post-hoc test, F (2, 77) = 0.7219, P=0.4891; Control n= 26, 15q13.3 HET Carrier n=26, 15q13.3 HET Proband n=28)
**(g)** Membrane resistance (Kruskal-Wallis test with Dunn's post-hoc test; Kruskal-Wallis statistic= 2.392, p=0.3024; Control n= 23, 15q13.3 HET Carrier n=24, 15q13.3 HET Proband n=23)
**(h)** Action potential amplitude from threshold (One-Way ANOVA with Tukey's post-hoc test, F (2, 67) = 0.7728, P=0.4658; Control n= 24, 15q13.3 HET Carrier n=24, 15q13.3 HET Proband n=23)
**(i)** Action potential width (Kruskal-Wallis test with Dunn's post-hoc test; Kruskal-Wallis statistic=3.201, p=0.2018; Control n= 24, 15q13.3 HET Carrier n=23, 15q13.3 HET Proband n=23)
**(j)** Action potential half-width (Kruskal-Wallis test with Dunn's post-hoc test; Kruskal-Wallis statistic=2.734, p=0.2548; Control n= 24, 15q13.3 HET Carrier n=23, 15q13.3 HET Proband n=23)
**(k-o)** Intrinsic electrophysiological properties of Family 3 iNeurons.
**(k)** Resting membrane potential (Unpaired t-test two-tailed, t=0.6603, df=56; Control n=30, 15q13.3 HET n=28)
**(l)** Membrane resistance (Mann-Whitney test two-tailed exact, U=340, p-value =0.4060; Control: 400.5, n=27, 15q13.3 HET: 417.2, n=29).
**(m)** Action potential amplitude from threshold (Unpaired t-test two-tailed, p=0.1341, t=1.520, df=57; Control n=30 neurons, 15q13.3 HET n=29).
**(n)** Action potential width (Unpaired t-test two-tailed, p=0.5571, t=0.5907, df=56; Control: n=30 neurons, 15q13.3 HET: n=28 neurons).
**(o)** Action potential half-width (unpaired t-test two-tailed, p=0.3496, t=0.9431, df=57; Control: n=30 neurons, 15q13.3 HET: n=29 neurons).

#### Supplementary Figure 10

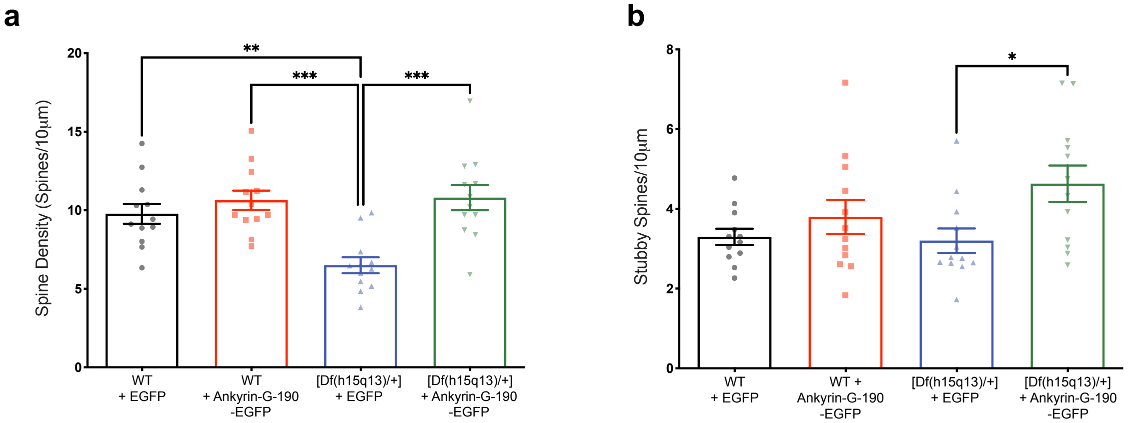


**Supplementary Figure 10. Additional dendritic spine analysis from WT and *Df(h15q13)/+* cortical neurons expressing Ankyrin-G-190-EGFP**

**(a)** Expression of Ankyrin-G-190-EGFP in *Df(h15q13)/+* neurons increases spine density to WT levels. n=12 neurons per condition from 3 mouse cultures. **p<0.01, ***p<0.001; One-Way ANOVA with Tukey's post-hoc test; F (3, 44) = 9.566, P<0.0001.
**(b)** Expression of Ankyrin-G-190-EGFP in *Df(h15q13)/+* neurons increases stubby spine density. n=12 neurons per condition from 3 mouse cultures *p<0.05; One-Way ANOVA with Tukey's post-hoc test; F (3, 44) = 3.234, P=0.0311.
